# Disrupted circadian core-clock oscillations in Type 2 Diabetes are linked to altered rhythmic mitochondrial metabolism

**DOI:** 10.1101/2021.02.24.432683

**Authors:** Brendan M. Gabriel, Ali Altıntaş, Jonathon A.B. Smith, Laura Sardon-Puig, Xiping Zhang, Astrid L. Basse, Rhianna C. Laker, Hui Gao, Zhengye Liu, Lucile Dollet, Jonas T. Treebak, Antonio Zorzano, Zhiguang Huo, Mikael Rydén, Johanna T. Lanner, Karyn A. Esser, Romain Barrès, Nicolas J. Pillon, Anna Krook, Juleen R. Zierath

## Abstract

Circadian rhythms are generated by an auto-regulatory feedback loop composed of transcriptional activators and repressors. Disruption of circadian rhythms contributes to Type 2 diabetes (T2D) pathogenesis. We elucidated whether altered circadian rhythmicity of clock genes is associated with metabolic dysfunction in T2D. Transcriptional cycling of core clock genes *ARNTL, CLOCK*, *CRY1* and *NR1D1* was altered in skeletal muscle from individuals with T2D and this was coupled with reduced number and amplitude of cycling genes and disturbed circadian oxygen consumption. Mitochondrial associated genes were enriched for differential circadian amplitudes in T2D, and positively correlated with insulin sensitivity. ChIP-sequencing identified CLOCK and BMAL1 binding to circadian mitochondrial genes associated with insulin sensitivity, implicating regulation by the core clock. Mitochondria disruption altered core-clock gene expression and free-radical production, phenomena that were restored by resveratrol treatment. We identify bi-directional communication between mitochondrial function and rhythmic gene expression, processes which are disturbed in diabetes.

## Introduction

Type 2 diabetes (T2D) is a growing global health problem, with skeletal muscle insulin resistance a primary defect in the pathology of this disease. While the etiology of this disease is complex, perturbed circadian rhythms from shift-work, sleep disorders, and social jet-lag are associated with obesity, T2D, and related co-morbidities (Mokhlesi et al., 2019; Shan et al., 2018; Tan et al., 2018; Vetter et al., 2018), highlighting the critical role of this circuit on metabolic health. Cell autonomous circadian rhythms are generated by a transcription-translation auto-regulatory feedback loop composed of transcriptional activators CLOCK and BMAL1 (*ARNTL*) and their target genes Period (*PER*), Cryptochrome (*CRY*), and REV-ERBα (*NR1D1*), which rhythmically accumulate and form a repressor complex that interacts with CLOCK and BMAL1 to inhibit transcription (Takahashi, 2015). Disruption of the molecular-clock in skeletal muscle leads to obesity and insulin resistance in mouse models (Dyar et al., 2014; Harfmann et al., 2016; Schiaffino et al., 2016). While disrupted circadian rhythms alter metabolism, the extent to which these processes are impaired in people with T2D is unknown.

Several lines of evidence suggest that the link between dysregulated molecular-clock activity and T2D or insulin resistance may be tissue-dependent. In white adipose tissue, core-clock (*PER1, PER2, PER3, CRY2, BMAL1, and DBP*), clock-related (*REVERBα*) and metabolic (*PGC1α*) genes showed no difference of rhythm and amplitude between individuals with normal weight, obesity, or T2D over a time-course experiment (Otway et al., 2011). Conversely, in human leucocytes collected over a time-course experiment, mRNA expression of *BMAL1, PER1, PER2* and *PER3* was lower in people with T2D as compared to non-diabetic individuals (Ando et al., 2009). Additionally, *BMAL1, PER1* and *PER3* mRNA expression in leucocytes collected from people with T2D is inversely correlated with HbA1c levels, suggesting an association of molecular-clock gene expression with T2D and insulin resistance. Furthermore, in pancreatic islets from individuals with T2D or healthy controls, *PER2, PER3* and *CRY2* mRNA expression is positively correlated with islet insulin content and plasma HbA1c levels (Stamenkovic et al., 2012). Thus, there may be tissue-specificity of molecular-clock regulation, which contributes to clinical outcomes related to insulin sensitivity and T2D etiology. The underlying mechanisms regulating metabolic rhythmicity and, particularly, whether rhythmicity is lost in T2D remain incompletely understood.

At the cellular level, primary human myotubes maintain a circadian rhythm, with the amplitude of circadian gene -*REV-ERBα* correlating with the metabolic disease state of the donor groups (Hansen et al., 2016). This apparent link between the skeletal muscle molecular-clock and insulin sensitivity may be partly mediated by molecular-clock regulation of metabolic targets. ChIP-sequencing has revealed distinct skeletal muscle-specific BMAL1 and REV-ERBα cistromes (Dyar et al., 2018), with prominent molecular-clock targeted pathways including mitochondrial function, and glucose/lipid/protein metabolism (Dyar et al., 2018; Pastore and Hood, 2013). Moreover, these metabolic pathways may participate in retrograde signaling to control aspects of the molecular-clock. Pharmacological inhibition of *DRP1*, a key regulator of mitochondrial fission and metabolism, alters the period length of BMAL1 transcriptional activity in human fibroblasts (Schmitt et al., 2018). However, the signals and the clock-derived alterations that govern the rhythmicity of metabolism remain incompletely understood. Despite the growing evidence that several metabolic pathways are under circadian control, it is not clear whether circadian rhythmicity of the intrinsic molecular-clock is altered in T2D. Here we determined whether circadian control of gene expression and metabolism is altered at the cellular level in skeletal muscle from individuals with T2D.

## Results

### Intrinsically dysregulated circadian rhythm of gene expression in T2D myotubes

Skeletal muscle biopsies were obtained from men with either T2D or normal glucose tolerance (NGT) (Figure 1A). Primary myotubes cultures were prepared, synchronized and harvested every 6 hours over 42 hours and the transcriptome was analyzed by RNA-sequencing (RNA-Seq). To determine whether genes displayed a rhythmic cycle over ∼24 hours, we analyzed the RNA-seq expression values using “rhythmicity analysis incorporating non-parametric methods” (RAIN) algorithm with longitudinal method (Thaben and Westermark, 2014) (Supplemental Table 1). Myotubes from T2D donors displayed fewer cycling rhythmic genes as compared to myotubes from NGT donors and the rhythmic genes in NGT were significantly associated with the rhythmic genes in T2D (Figure 1B, Fisher exact test, p=3.91e- 58). To induce acute insulin resistance and partly mimic a T2D milieu (Vlavcheski et al., 2020), myotubes from NGT and T2D donors were treated with high glucose and insulin (50 nM insulin, 25 mM glucose) for 24 hours prior to serum shock. High concentration glucose and insulin treatment reduced the number of cycling genes in myotubes from both T2D and NGT donors (Figure 1C). We performed gene set enrichment analysis (GSEA) and found that reactome pathways (Jassal et al., 2020) (Figure 1D) that were enriched for cycling genes in the myotubes from the NGT donors included metabolic pathways such as ‘*Regulation of lipid metabolism by PPARA’* and ‘*PPARA activates gene expression*’, while the reactome pathway ‘*Circadian Clock’* was enriched in myotubes from the T2D donors. No reactome pathways were enriched in NGT (high glucose and insulin concentration), while ‘Circadian Clock’ was enriched in T2D (high glucose and insulin concentration). These data indicate that there are more metabolic genes cycling in cells from NGT donors. To assess the altered rhythmicity, so called differential rhythmicity, between NGT and T2D, the RNA-seq expression values from the NGT and T2D donors were compared to each other using DODR algorithm (Thaben and Westermark, 2014). *BMAL1 (a.k.a. ARNTL)*was the top differentially rhythmic gene (3 total, Supplemental Table 2) between NGT and T2D donors (Figure 1E, FDRDODR=0.096). Several molecular-clock genes, as well as clock-output genes, were exclusively cycling in either NGT or T2D, including *CLOCK*, *CRY1* and, *NR1D1* (FDRRAIN<0.1) (Figure 1E). These results indicate that myotubes from the NGT donors had more genes displaying rhythmic behavior. Moreover, cycling genes in NGT cells were enriched in metabolic pathways as compared to T2D cells, as well as cells treated with a high concentration of glucose and insulin. Additionally, several core clock genes displayed differential rhythmicity (DODR) or were exclusively rhythmic (RAIN) in myotubes from NGT and T2D donors.

**Figure 1.**
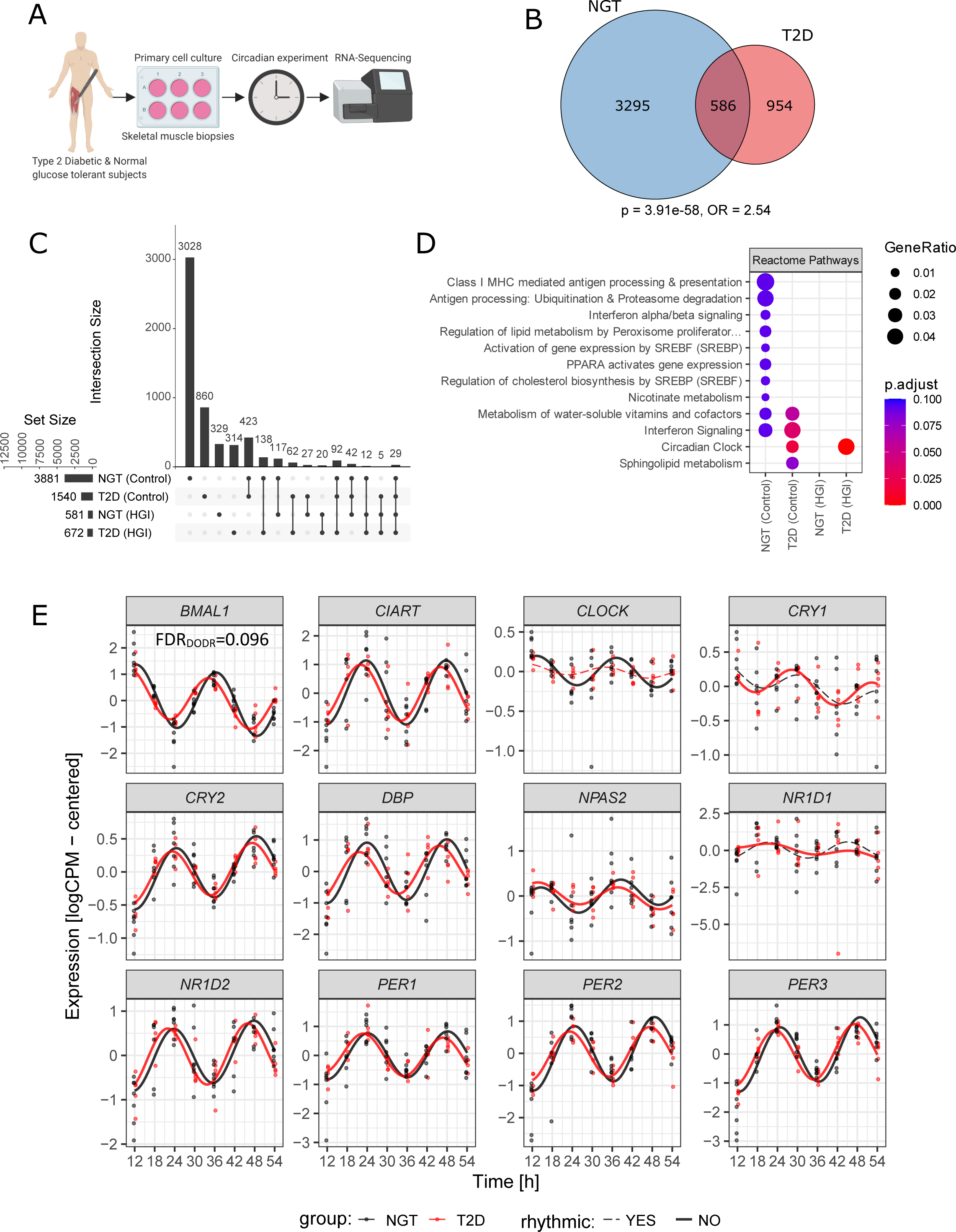
Intrinsically dysregulated circadian rhythm of gene expression in T2D. (A) Schematic overview of primary cell culture circadian experiment and RNA-Sequencing. RNA-seq of primary human skeletal muscle cells from men with NGT (n=7) or T2D (n=5). (B) Venn diagram showing the overlapping rhythmic genes between NGT and T2D. Overlapping rhythmic genes were significantly enriched (Fisher’s exact test, p = 3.91e-58; Odds-ratio, OR = 2.54, background = 18,483). (C) Number of circadian genes identified via RAIN analysis (FDR<0.10). (D) Circadian gene enrichment results using over-representation analysis (Fisher’s exact test) and Reactome pathways. Circadian genes identified via RAIN analysis (FDR<0.10) and top 10 enriched Reactome pathways are shown (FDR<0.10). (E) Circadian rhythmicity of core clock genes. Red=T2D, Black=NGT. Lines show the harmonic regression fits and solid line indicates circadian (FDR<0.10) genes while dashed lines indicate non-circadian genes. Time points are hours post-synchronization.

### Altered peak-time signature of cycling genes in myotubes from T2D donors

Analyzing the same RNA-sequencing data described in Figure 1, we used the RAIN algorithm to determine peak time of cycling genes in the myotubes. T2D displayed an altered pattern of cycling gene peaks as compared to NGT (Figure 2A and 2B). In percentage terms, T2D had the highest number of cycling genes displaying peaks at 24 hours, whereas NGT had the highest number of cycling genes displaying peaks at 12 hours. The number of cycling genes at each phase peak was more similar between T2D and NGT when cells were treated with a high concentration of glucose and insulin (Figure 2C and 2D). However, the peak at which both NGT and T2D had the largest amount of cycling genes remained consistent within diagnosis groups, with the highest number of cycling genes displaying peaks at 12 hours (NGT) and 24 hours (T2D), respectively. When considering the molecular-clock genes and clock-output genes (Figure 2E), *NPAS2, DBP* and *NR1D1* displayed different peak times between NGT and T2D. Additionally, *NR1D1* and *PER3* displayed different peak times between myotubes from NGT and NGT donors treated with a high concentration of glucose and insulin. We then performed a gene enrichment analysis (ORA) for each condition and gene peak time. Several metabolic pathways displayed enrichment for cycling genes at peak times 6 and 12 hours in NGT cells, but not in any other condition (Figure 2F). For example, the pathway ‘*Metabolism of carbohydrates’* was enriched at 6 hours only in NGT, whereas ‘*Regulation of lipid metabolism by PPARA’* was enriched at 12 hours, in NGT alone. Thus, the myotubes from the NGT and T2D donors have an altered peak-time gene expression signature that is somewhat conserved even when cells are treated with a high concentration of glucose and insulin.

**Figure 2.**
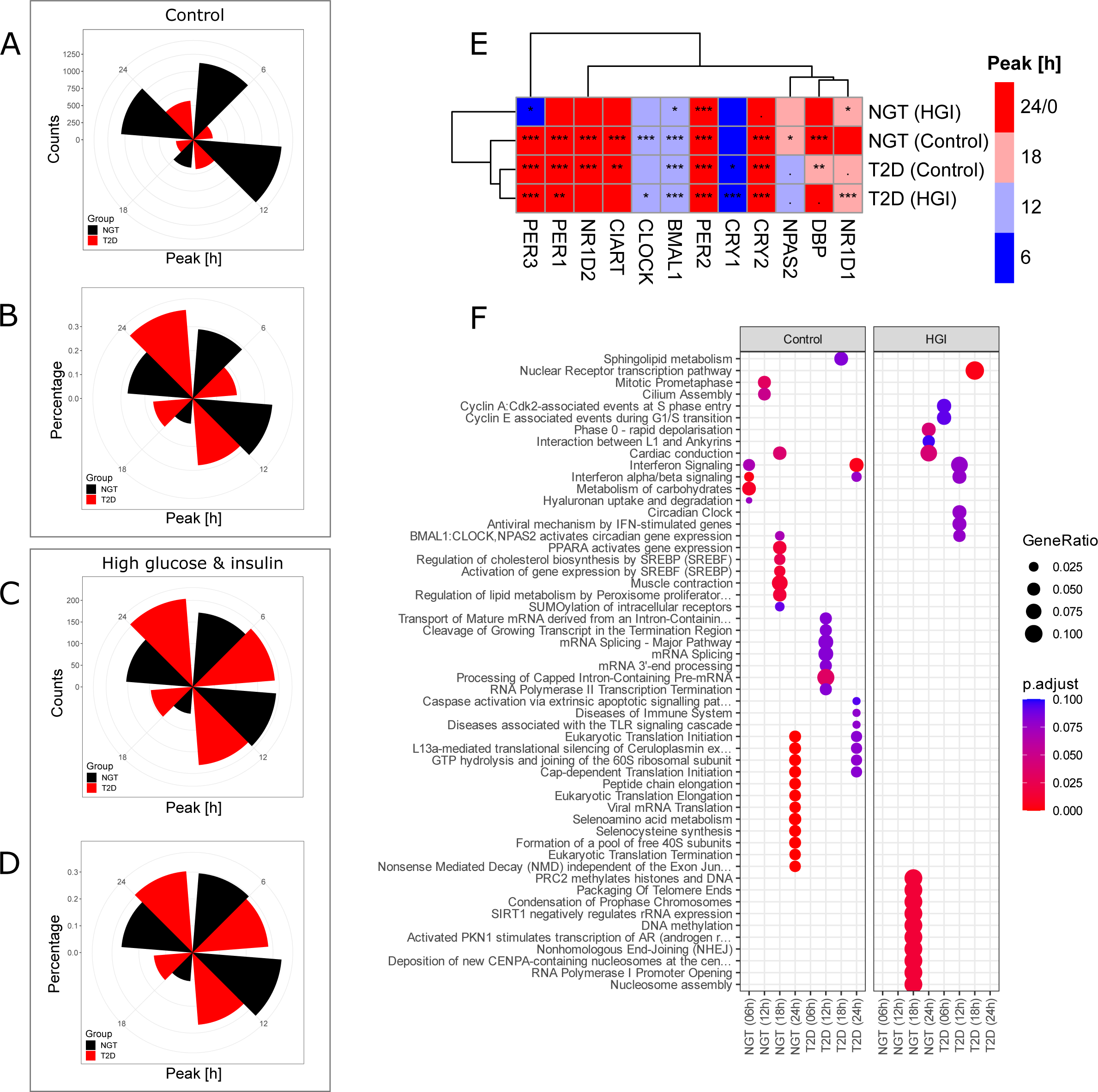
Altered peak-time signature of cycling genes in T2D. (A) Number of circadian genes at each peak time for control treatments. (B) Percentage of circadian genes at each peak time for control treatments. (C) Number of circadian genes at each peak time when treated with high concentrations of glucose and insulin. (D) Percentage of circadian genes at each peak time when treated with high concentrations of glucose and insulin. (E) Heatmap showing the peak times of core clock genes for control conditions and cells treated with high concentrations of glucose and insulin. Colors represent the peak times. Hierarchical clustering was performed by using geodesic distance and ‘ward.D2’ algorithm. Star signs represent the adjusted p-values from rhythmicity analysis (RAIN): “***” (p < 0.001), “**” (0.001 - 0.01), “*” (0.01 - 0.05), “.” (0.05 – 0.1) and empty boxes (0.1 – 1). (F) Reactome pathways enriched at each time point in NGT (control), T2D (control), NGT (high concentration of glucose and insulin.), and T2D (high concentration of glucose and insulin.).

### Reduced amplitudes of rhythmic mitochondrial, and overall gene expression and ablated rhythmic mitochondrial metabolism in T2D

An important factor which is often physiologically relevant in circadian and diurnal biology is the magnitude of cycling peaks and nadirs over the course of a cycle, which can be quantified by measuring the amplitude of cycling patterns. We used harmonic regression to determine relative amplitude of cycling gene expression in myotubes from NGT and T2D donors within the RNA-sequencing data-set. The mean log2 amplitude of cycling genes was lower in T2D as compared to NGT (Figure 3A; p=2.2e-16, two-sided Kolmogorov-Smirnov test). To determine if there was a specific cellular location that was enriched for amplitude divergence, we next performed GSEA using ranked log2-fold-changes of relative amplitudes. This analysis determined that the enriched Gene Ontology (GO) cellular components for reduced amplitudes in T2D compared to NGT were‘*Ficolin-1-rich granule lumen’*, *‘Clathrin vesicle coat’*, *‘Organelle inner membrane’*, and *‘Mitochondrial inner membrane’, ‘Organelle Envelope’* and ‘*Mitochondrial Membrane’* (Figure 3B).

**Figure 3.**
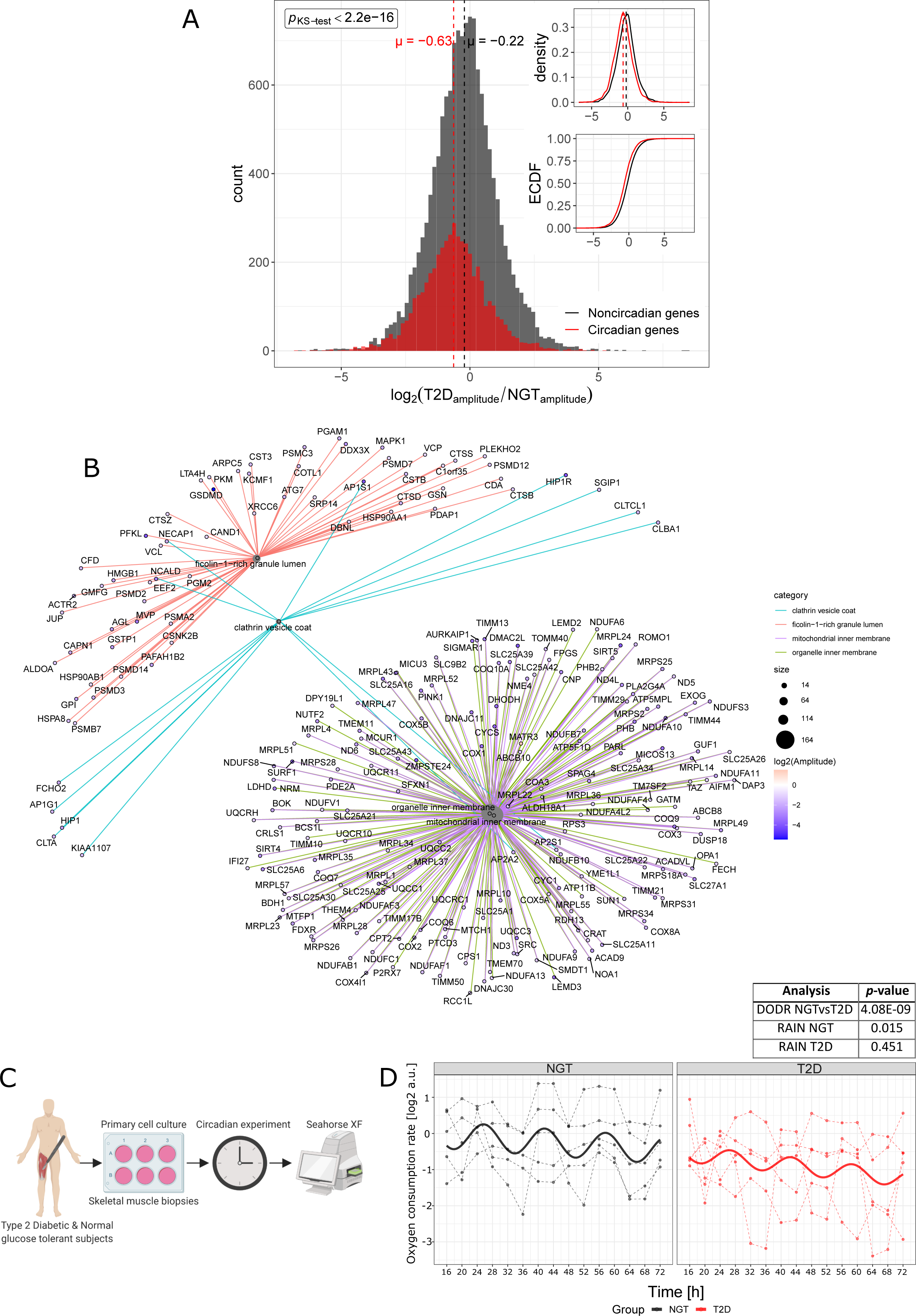
Reduced amplitudes of rhythmic mitochondrial, and overall gene expression and ablated rhythmic mitochondrial metabolism in T2D. (A) Log2 relative amplitude of circadian genes for T2D as compared to NGT, relative amplitudes determined via harmonic regression with first degree polynomial trend using mean centered data. Histogram of circadian genes (red) and all genes (black), log2 relative amplitude of T2D vs NGT. The main panel shows count histogram of log2 relative amplitudes of circadian genes (red) and noncircadian genes (black). Panel on right top show the density distributions and ECDF (Empirical Cumulative Distribution Function). Red=circadian, determined via RAIN algorithm (FDR<0.1), Black=non-circadian. Log2 relative amplitude distributions were significantly different in circadian genes compared to noncircadian genes (*p*=1.28e-6, two-sided Kolmogorov-Smirnov test). µ=population mean. (B) GO term – Gene interaction plot of top enriched cellular component for ranked log2 relative amplitude changes of GO terms *mitochondrion, mitochondrial membrane, organelle envelope*. (C) Schematic of circadian basal cellular oxygen consumption rate time-course experiment. (D) Relative oxygen consumption rate of synchronized myotube cultures from donors with NGT (black) versus T2D (red), as measured by Seahorse XF Analyzer (Agilent) for n=5 individuals in both groups. Differential rhythmicity (DODR, period = 16 hours) and rhythmicity (RAIN) analysis statistics are shown in table inset. See also Figure S1.

Given that reduced amplitude of a mitochondrial GO cellular component was observed in myotubes from the T2D donors, and that T2D is associated with reduced skeletal muscle mitochondrial function and metabolic inflexibility (Hesselink et al., 2016), we investigated whether synchronized myotubes from NGT and T2D donors displayed circadian oscillations of oxygen consumption rate (OCR; a proxy for mitochondrial metabolism, Figure 3C). We observed differential rhythmicity of basal OCR between NGT and T2D (DODR, FDR=4.08e-09) (Figure 3D). Additionally, NGT displayed cycling OCR with a 16-hour period (FDRRAIN=0.015) whereas T2D did not (nor any other period). Additionally, we performed a separate MitoStress test at a single timepoint (Zeitgeber Time: ZT24) after serum shock (Figure S1). In this assay, there were no differences in mitochondrial function between the myotubes from NGT and T2D donors. Thus, the difference of rhythmic mitochondrial function between the myotubes from the NGT and T2D donors may be derived from temporal regulation of basal oxygen consumption, rather than differences in mitochondrial function *per se*. Overall, the data in Figure 3 demonstrate that myotubes from T2D donors have reduced mitochondrial metabolic rhythm in terms of both oxygen consumption rate and mitochondrial related gene amplitudes.

### The mitochondrial inner-membrane is enriched for lower circadian amplitudes and expression of genes that correlate with whole-body insulin sensitivity in T2D

To test the *in vivo* clinical relevance of our findings in the primary myotube cultures, we performed a transcriptomic analysis on *vastus lateralis* muscle biopsies obtained from men with NGT (n=22) or T2D (n=22) at a single timepoint (Figure 4A). Additionally, a hyperinsulinemic-euglycemic clamp was performed to determine whole-body insulin sensitivity (M-value, analysis pipeline shown in Figure 4A). As expected, insulin sensitivity was greater in men with NGT as compared to T2D (Figure 4B, p<0.0001). The relationship between basal skeletal muscle gene expression and insulin sensitivity across the whole cohort was assessed by Spearman’s rank correlation (Figure 4A). A gene-set enrichment analysis (GSEA) with ontology terms from GO cellular components was performed using Spearman’s correlation coefficient as ranking metric and found that most of the enriched GO cellular components were related to the mitochondria (FDR < 0.10, Figure 4C and 4D). We then compared the enriched cellular components from the gene set enrichment analysis of log2-fold-change relative amplitudes that were down-regulated in T2D (Figure 4C). Two cellular components were consistent between these two analyses *‘Mitochondrial inner membrane’,* and ‘*Organelle inner membrane’* (Figure 4D). Our analysis points at the inner-mitochondria as a cellular location implicated in both the regulation of insulin sensitivity and associated with impaired cycling behaviors in skeletal myotubes.

**Figure 4.**
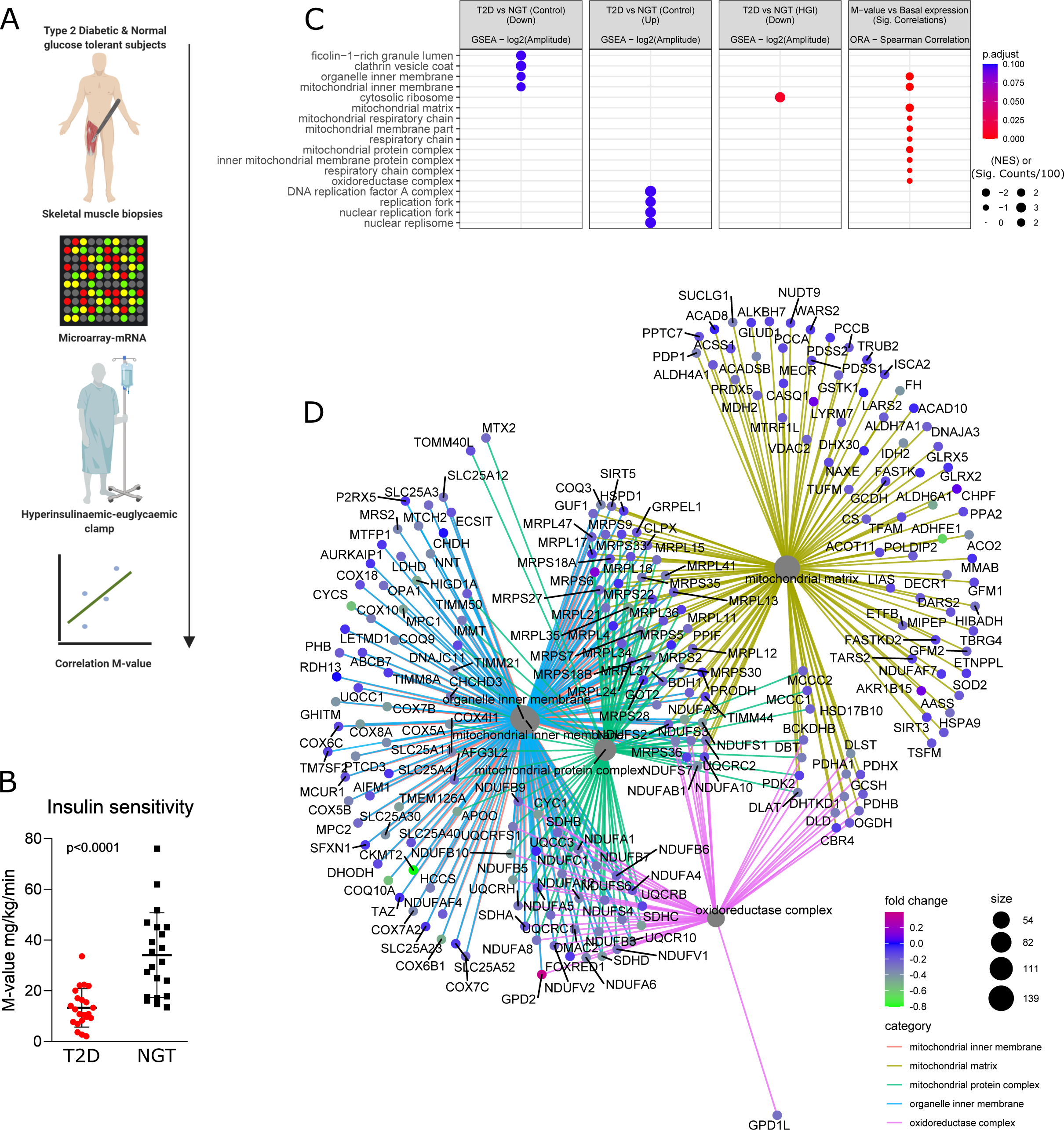
Mitochondrial pathways are enriched for lower T2D circadian amplitudes and genes that correlate with whole-body insulin sensitivity. (A) Schematic of experimental design. (B) Insulin sensitivity (M-value) of T2D and NGT participants. Student’s t-test. (C) Enrichment analysis based on microarray of skeletal muscle biopsies obtained in the fasted state from 22 men with NGT and 22 men with T2D obtained in the fasted state. Differentially enriched GO cellular components (GO:CC), ranked by FDR-values. Column 1 and 2: Top enriched cellular components for ranked log2 relative amplitude changes, Column 3: Pathway enrichment calculated from genes correlating FDR<0.10) with M-value (whole-body insulin sensitivity) *in vivo*. There were no enriched pathways in the T2D versus NGT (high concentration of glucose and insulin.) (Up) condition. (D) GO term – Gene interaction plot of GO:CC enrichments calculated from genes correlating (FDR*<*0.1) M-value (whole-body insulin sensitivity) *in vivo*: *organelle inner membrane, mitochondrial inner membrane, mitochondrial protein complex, mitochondrial matrix, oxidoreductase complex*.

### Mitochondrial pathways are enriched for circadian genes with Clock and Bmal1 binding, and are associated with whole-body insulin sensitivity

In the current study, we describe intrinsically dysregulated rhythm, peak time, and amplitudes of cycling genes in myotubes from T2D as compared to NGT donors. Reduced amplitudes of cycling gene expression in T2D were particularly associated with mitochondrial related genes, and this was coupled with functionally dysregulated mitochondrial rhythmic metabolism. However, the regulatory directionality of these phenomena (*i.e.* rhythmic dysfunction of the molecular-clock and mitochondrial metabolism) is unclear. Whether both BMAL1 and CLOCK partially regulate mitochondrial metabolism in skeletal muscle is also unclear. We found that these molecular-clock genes were rhythmically dysregulated in myotubes from T2D as compared to NGT donors. Therefore, we tested whether BMAL1 and CLOCK binding was associated with mitochondrial genes involved in insulin sensitivity and dysregulated mitochondrial metabolic rhythms, which would indicate that dysregulated BMAL1 and CLOCK lead to disrupted mitochondrial rhythms in skeletal muscle of people with T2D. We performed ChIP-sequencing of mouse skeletal muscle using BMAL1 and CLOCK antibodies and assessed peaks closest to a gene’s transcription start site. Firstly, we assessed whether circadian genes identified in myotubes from NGT and T2D donors were associated with BMAL1 and CLOCK binding to their counterpart genes (Figure 5A). Out of all the myotube conditions, only NGT was associated with BMAL1 and CLOCK binding, while T2D cells treated with a high concentration of glucose and insulin were associated with CLOCK binding exclusively. We then performed integrative clustering analysis between the ChIP-sequencing data, skeletal muscle gene expression correlated with insulin sensitivity, and circadian genes identified within myotubes from NGT and T2D donors incubated in the absence or presence of a high concentration of glucose and insulin. (Figure 5B). To integrate all different experimental outcomes, the data was presented as a binary matrix while 1 shows the presence and 0 shows the absence of rhythmicity, correlation or ChIP-seq peaks. The integrated data was subjected to clustering analysis, which resulted in 10 distinct clusters (Figure 5C). Clusters associated with BMAL1 and CLOCK binding were also associated with circadian rhythm in NGT (clusters 6, 7, and 8) and T2D (clusters 1, 2, 3, and 4). Several clusters demonstrated an association between T2D control conditions and T2D (high concentration of glucose and insulin) circadian genes, while NGT (high concentration of glucose and insulin)circadian genes clustered with very few other categories.

**Figure 5.**
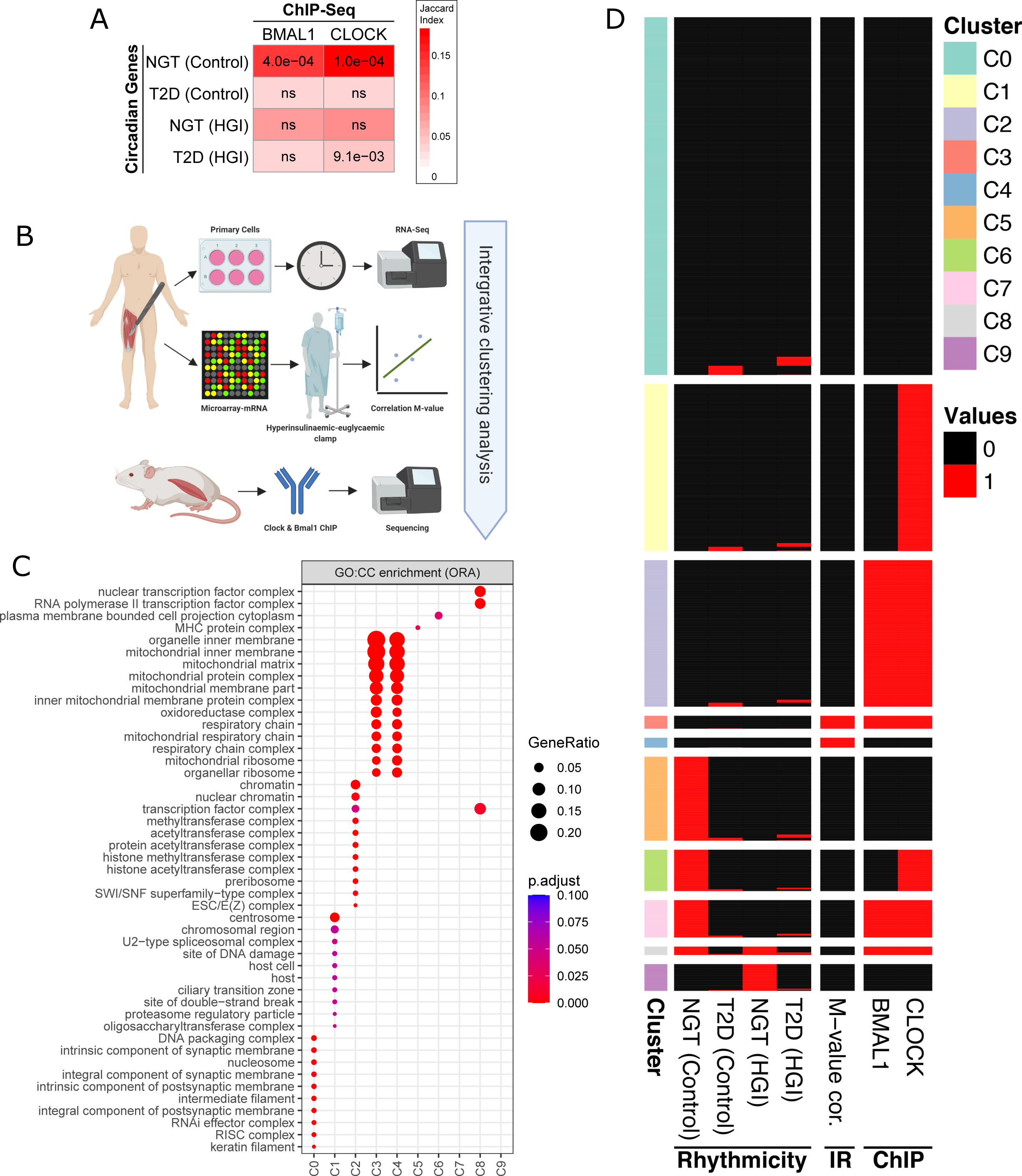
Mitochondrial pathways are enriched for circadian genes with Clock and Bmal1 binding, and are associated with whole-body insulin sensitivity. (A) Enrichment analysis between circadian genes and ChIP-seq peaks. The association between circadian genes in each disease-treatment group and BMAL1/CLOCK bound genes were tested using Fisher’s exact test and p-values are adjusted for multiple testing with Benjamini-Hochberg method. Columns: BMAL1/CLOCK ChIP-seq experiments; rows: Rhythmic genes in NGT and T2D for both control conditions and high concentration of glucose and insulin treatment groups; numbers: adjusted p-values (ns: not significant); colors: Jaccard similarity index indicating the percentage of genes overlapping in each dataset. (B) Schematic of experimental design used for the integrative analysis. (C) Heatmap showing the clusters from the integrative analysis. Binary data used for clustering with “clust” algorithm (Abu-Jamous and Kelly, 2018) and resulted in 10 clusters. Columns “NGT (Control)”, “T2D (Control)”, NGT (high concentration of glucose and insulin.)”, “T2D (high concentration of glucose and insulin.)” represents circadian (black) and noncircadian (white) genes; “M-value cor” shows whether a significant (black) correlation between the insulin sensitivity metric (M-value) and basal gene expression; “BMAL1” and “CLOCK” show whether a ChIP-seq peak for the given protein was detected (black) or not (white). (D) Gene enrichment analysis using over-representation (Fisher’s exact test) with GO cellular components was performed for each cluster identified in (B).

Genes that correlated with insulin sensitivity formed a cluster with BMAL1 and CLOCK binding, and circadian genes within myotubes from T2D donors (including when treated with a high concentration of glucose and insulin)(Cluster 3, and 4). We then performed gene set enrichment analysis of GO cellular components (Figure 5D). Of note, clusters 3 and 4 were enriched for genes associated mainly with the inner-mitochondria. There appeared to be few clustering events between insulin sensitivity-associated genes, genes with BMAL1 and CLOCK binding, and genes that were circadian in NGT. Thus, mitochondrial rhythmic metabolism within NGT may be partly independent of genes with CLOCK or BMAL1 binding.

### Pharmacological, genetic, and siRNA-mediated disruption of inner-mitochondrial metabolism results in altered clock-gene expression

Our data provide evidence that mitochondrial metabolic rhythms in skeletal muscle from individuals with NGT may be partly independent of direct core-clock control. Additionally, internal mitochondrial gene expression was associated with whole-body insulin sensitivity, inner-mitochondrial genes also had reduced amplitudes in myotubes from T2D donors. Targets involved in internal mitochondrial functionality can act as regulators of molecular-clock expression and function (Lassiter et al., 2018; Schmitt et al., 2018). Thus, we hypothesized that internal mitochondrial functionality may play a role in regulating metabolic and molecular clock rhythms. To test this in myotubes from NGT donors, we used different compounds that target mitochondria, namely Carbonyl cyanide 4-(trifluoromethoxy)phenylhydrazone (FCCP), Oligomycin, and a mixture of Rotenone and Antimycin A (Rot/AA). After serum shock we incubated myotubes with these compounds for 4h (ZT14-ZT18). We measured mRNA expression of molecular clock-associated genes *DBP* (Figure 6A) and *NR1D1* (Figure 6B), and found that 0.33 µM of Rot/AA, and 2 µM of FCCP increased mRNA expression of *DBP* as compared to vehicle control-treated myotubes. Additionally, 2 µM of FCCP, and 1 µM of oligomycin increased mRNA expression of *NR1D1* as compared to vehicle control-treated myotubes. These data provide evidence that manipulation of internal mitochondrial function may play a role in altering clock-associated gene expression in primary human myotubes. However, compounds such as these may have off-target effects, and thus we tested this hypothesis in other models.

**Figure 6.**
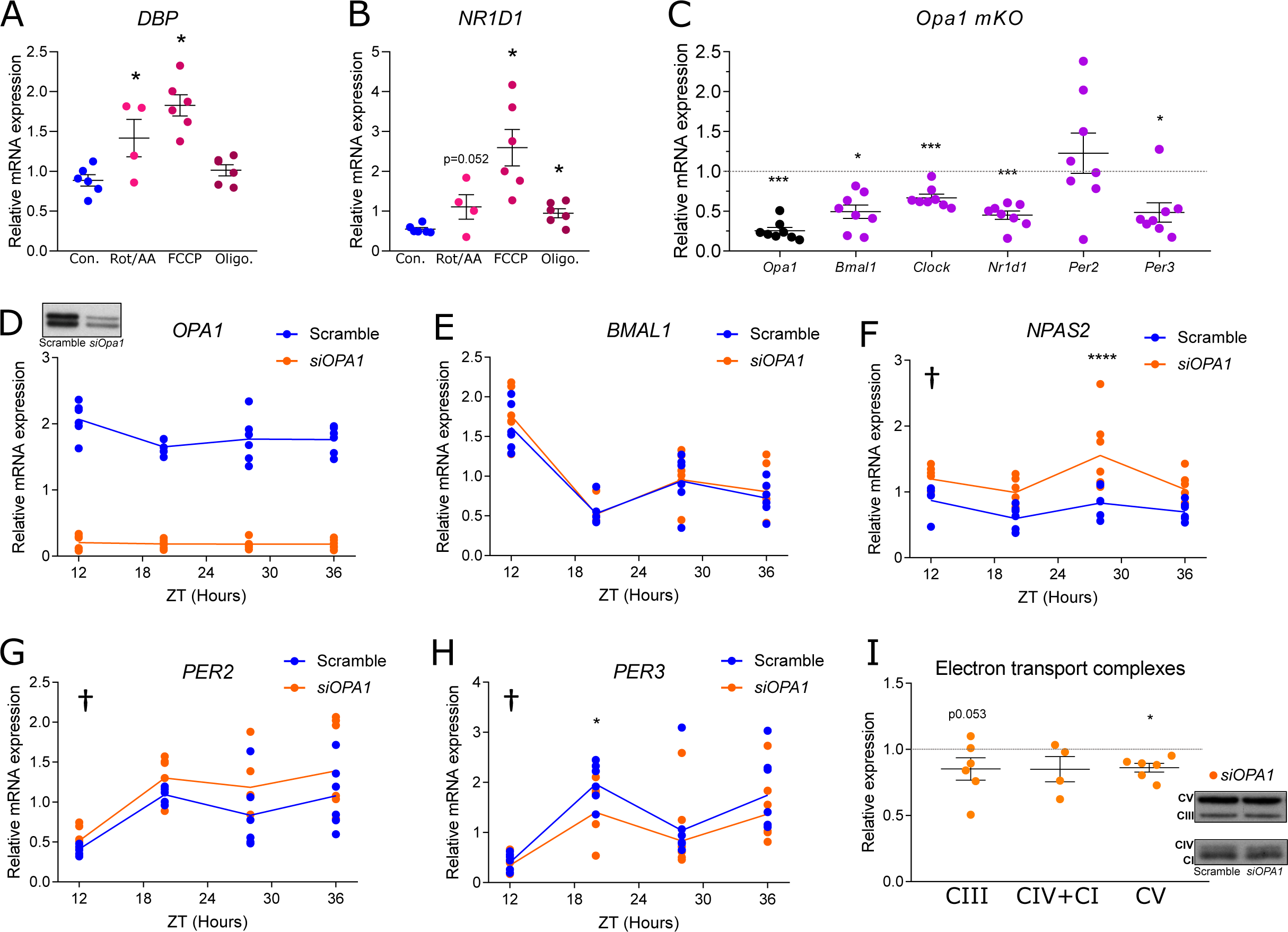
Pharmacological, genetic, and siRNA-mediated disruption of inner-mitochondrial metabolism results in altered clock-gene expression. (A) mRNA expression of molecular clock-associated gene DBP in myotubes from NGT donors (n=4-6 donors). One-way ANOVA, *=p<0.05 compared to vehicle control (Con.). (B) mRNA expression of molecular clock-associated gene NR1D1 in myotubes from NGT donors (n=4-6 donors). One-way ANOVA, *=p<0.05 compared to vehicle control (Con.). (C) *OPA1* and Clock gene expression (*Bmal1, Clock, Nr1d1, Per2, Per3*) from *Opa1 MKO- Opa1* skeletal muscle *Opa1^-/-^* mice (n=8) relative to control (n=7). Black=siRNA-target *Opa1* gene expression, purple=core-clock/associated gene expression One-way ANOVA, ***=*p*<0.001, *=*p*<0.5. (D) *OPA1 mRNA* expression, and protein abundance (inset) over time-course experiment after synchronization in primary human skeletal muscle cells treated with *siRNA* targeting *OPA1* (*siOPA1*) (n=6) compared to scramble *siRNA* (n=6). (E-H) Primary human skeletal muscle cells treated with siRNA targeting *OPA1* (n=6) (*siOPA1*). Molecular-clock genes *BMAL1* (E)*, NPAS2* (F), *PER2* (G), and *PER3* (H). †*p*<0.05 overall difference *siOPA1 vs*. scramble. \*\*\*\**p*<0.0001, \**p*<0.05 *siOPA1 vs*. Scramble at time point (2-way ANOVA). Genes are shown in synchronized cells and presented at each zeitgeber-time post serum shock. (I) Protein abundance of electron transport complexes. Orange siRNA targeting *OPA1*. One-way ANOVA, †=Overall effect of siOPA1 (*p*<0.05), \**p*<0.05 effect of siOPA1.

We sought to manipulate a target that both regulated inner-mitochondrial function in myotubes and displayed diurnal behavior *in vivo* in skeletal muscle biopsies from individuals with NGT. Meeting these criteria, Optic Atrophy Protein 1 (OPA1) is an established regulator of inner-mitochondrial morphology and function in skeletal muscle (Lodi et al., 2011; Rodriguez-Nuevo et al., 2018; Sebastian et al., 2017; Tezze et al., 2017), and displays a day-night rhythm of protein abundance in skeletal muscle from healthy individuals, alongside oscillating mitochondrial function (van Moorsel et al., 2016). *OPA1* did not display cycling mRNA in myotubes from NGT or T2D donors when adjusted for multiple comparisons. However, *OMA1* and *HIGD2A* are responsible for the cleavage and stabilization of OPA1 protein (Anand et al., 2014; Salazar et al., 2019), respectively, and both of these genes displayed cycling mRNA patterns in myotube cultures from NGT, but not T2D donors (as determined via RAIN, Supplementary Figure 3, and Supplementary Table 1), suggesting circadian regulation of OPA1 protein. *OPA1* basal gene expression also correlated with insulin sensitivity in the microarray data (Spearman’s correlation, FDR=.002). To further elucidate the effect of mitochondrial disruption in relation to the molecular-clock machinery, we performed investigations in a skeletal muscle-specific *Opa1^-/-^* mouse model. Molecular-clock genes *Clock, Bmal1, Nr1d1* and *Per3* (Figure 6C) were decreased in *Opa1^-/-^* mice, suggesting retrograde signaling from OPA1 to the molecular-clock.

However, the *Opa1^-/-^* mice we studied have a severe myopathic phenotype (Rodriguez-Nuevo et al., 2018), we thus aimed to recapitulate this finding in myotube cultures from NGT donors (Figure 6D-6H). In these synchronized primary human skeletal myotubes, siRNA-targeted reduction of *OPA1* (Figure 6D) resulted in unchanged BMAL1 (Figure 6E) expression, while*NPAS2* (a paralogue and substitute of *Clock* (DeBruyne et al., 2007)) (Figure 6F), *PER2* (Figure 6G), and *PER3* (Figure 6H) all had altered *mRNA* expression in the depleted OPA1 condition. *OPA1* silencing also perturbed expression of complex V of the electron-transport chain (Figure 6 I). Collectively, these data suggest that manipulating inner-mitochondrial metabolism results in altered mRNA expression of clock genes.

### siRNA depletion of OPA1 increases mitochondrial reactive oxygen species, and OPA1- mediated changes in clock-gene expression can be restored by resveratrol treatment

Our data suggest that the inner-mitochondrion is a modulator of the molecular-clock genes *NPAS2*, *PER2* and *PER3* in human skeletal muscle (Figure 6), implicating retrograde signaling between OPA1 and the molecular-clock. This signaling pathway may also be relevant for insulin sensitivity, since myotubes from T2D donors have downregulated circadian amplitudes of mitochondrial genes, ablated rhythmic metabolism, and altered core-clock rhythm. There are at least two plausible candidate pathways through which inner-mitochondrial-mediated signaling to the core-clock may occur (Figure 7A). Firstly, NAD^+^ is a cofactor reflective of mitochondrial metabolism and NAD^+^-dependent sirtuins are regulators of molecular-clock genes (Peek et al., 2013). Secondly, oxygen sensing plays a key role in molecular signaling from the core-clock to mitochondria and vice-versa in skeletal muscle, via Hypoxia inducible factor-1 (HIF1-α) (Peek et al., 2017). Interestingly, HIF1-α mRNA displayed circadian rhythms in the myotube cultures from the NGT but not the T2D donors in our RNA-sequencing data (determined via RAIN, Figure 7B). HIF1-α expression is driven by hypoxia in skeletal muscle, and regulated by reactive oxygen species (ROS) (Hoppeler et al., 2003). Thus, we explored the circadian mRNA patterns within our RNA-sequencing data of myotubes from T2D and NGT donors to identify genes that localize to mitochondria, which are associated with ROS processing and NAD^+^/NADH metabolism. Genes that were circadian in either T2D or NGT are highlighted (Figure 7C, with several antioxidant enzymes showing circadian regulation in NGT but not T2D (*GPX1, GPX3, GPX4*). Additionally, several genes involved in NAD^+^/NADH were circadian in NGT but not T2D (*NARPT, NAMPT, NADSYN1*), while *NNT* was circadian in T2D but not NGT. Of note, *SOD2* (FDR=0.0005), *GPX1* (FDR=0.03), and *NAMPT* (FDR=0.007) correlated with insulin sensitivity in our microarray data (Spearman’s correlation). Given the apparent rhythmic differences in these metabolic pathways, we next assessed the corresponding metabolites in response to internal mitochondrion dysregulation. Control cells from NGT donors had differences in NAD^+^ concentration over time, while cultures treated with siRNA targeting OPA1 had an NAD^+^ concentration that did not differ over time However, there was no difference in NAD^+^ concentration between siOPA1 and SCR Figure 4D). We then assessed mitochondrial ROS concentration using live-cell microscopy and measured MitoSOX fluorescence, indicative of mitochondrial ROS. OPA1-depleted myotubes had increased MitoSOX fluorescence when perfused with Tyrode’s solution, as compared to scramble control (Figure 7E). We also treated OPA1-depleted myotubes with Resveratrol. Resveratrol can act as an antioxidant, and reduce HIF1-α activity, in addition to increasing SIRT1 activity (Repossi et al., 2020). Resveratrol reduced mitochondrial ROS levels in OPA1-depleted myotubes and rescued the OPA1-mediated changes in *NPAS2* expression (Figure 7F and G). Our results provide evidence to suggest that OPA1 regulation of mitochondrial ROS levels constitutes a feedback loop permitting bi-directional control of circadian metabolism in skeletal muscle.

**Figure 7.**
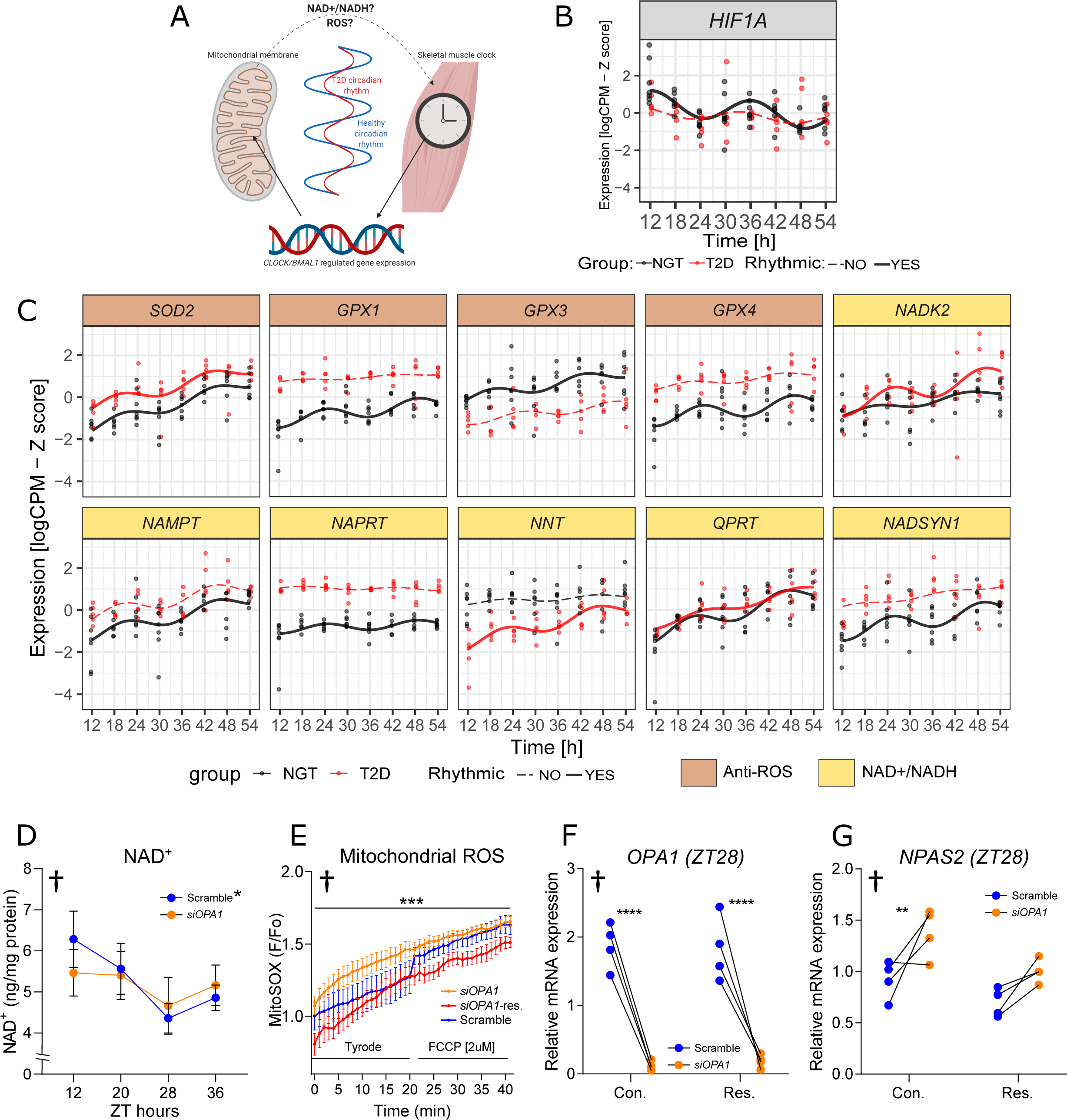
siRNA depletion of OPA1 increases mitochondrial reactive oxygen species, and OPA1-mediated changes in clock-gene expression can be restored by resveratrol. (A) Schematic diagram of hypothesized mechanism. (B) mRNA expression data from RNA sequencing of myotube cultures from donors with NGT or T2D (n=7, and 5, respectively) of HIF1α. Lines show the harmonic regression fits and solid line indicates circadian (adj.*p*<0.1) genes while dashed lines indicate non-circadian genes. Time points are hours post-synchronization. (C) mRNA expression data from RNA sequencing of myotube cultures from donors with NGT or T2D (n=7, and 5, respectively) of antioxidant enzymes, and those linked to NAD^+^/NADH metabolism that localize to mitochondria. Lines show the harmonic regression fits and solid line indicates circadian (adj.*p*<0.1) genes while dashed lines indicate non-circadian genes (Rain analysis). Time points are hours post-synchronization. (D) NAD^+^ concentration in synchronized primary human skeletal muscle cells (n=6) treated with either scramble (blue) or siRNA targeting *OPA1* (orange), two-way ANOVA, †overall variance of time, *<0.05 difference between timepoints (at 12 vs 20 hours, and 28 vs 36 hours) only in scramble condition. (E) Primary human skeletal muscle cells treated either with scramble-scramble siRNA or siOPA1-siRNA targeting *OPA1* (n=6), *OPA1* gene expression. live-cell microscopy measuring MitoSOX fluorescence, indicative of mitochondrial ROS. OPA1 depleted myotubes had increased MitoSOX fluorescence perfused with Tyrode’s solution, compared to scramble control. Resveratrol reduced ROS levels in cells with depleted OPA1 expression compared to siOPA1 and scramble control. Mixed effects-analysis, †=*p*<0.05 overall difference between conditions, ***=*p*<0.0001 difference between siOPA1 vs siOPA1-resv, Scramble vs siOPA1, and Scramble vs siOPA1-resv. (F and G) Gene expression of synchronized cells from men with NGT (n=4) were treated with either scramble (blue) or siRNA targeting *OPA1* (orange) at ZT28. Cells were either treated with 10 µM Resveratrol or vehicle (control) for the duration of the experiment, subsequent to synchronization. Relative to scramble siRNA at ZT28. Control conditions are presented as a dotted line in the figures. Two-way ANOVA, †=*p*<0.05 overall difference, \*\**p*<0.01 control-Scramble *vs*. Control-treated *siOPA1*. ***p<0.001 *siOPA1 vs* scramble. (F) *OPA1* and (G) *NPAS2* gene expression. Results are mean±SEM.

## Discussion

T2D shares several molecular and pathophysiological phenotypes with ageing (Cartee et al., 2016), which is associated with disruption of molecular-clock genes, in addition to dysregulation of other rhythmic biological processes (Zwighaft et al., 2015). Moreover, defective mitochondrial function and content is implicated in skeletal muscle insulin resistance and T2D pathogenesis (Ruegsegger et al., 2018; Schrauwen et al., 2010; Sebastian et al., 2017). Our data provide mechanistic insight into this metabolic dysfunction and show that myotubes from individuals with T2D have an intrinsically disrupted circadian rhythm, at the global transcriptomic and molecular-clock specific level. Furthermore, an acute treatment of myotubes with a high concentration of glucose and insulin, partially mimicking a diabetic milieu, somewhat replicates the reduced number of rhythmic genes in T2D. However, clear differences between T2D and treatment with high glucose and insulin concentrations exist, including the percentage of genes at each peak time, which were more similar within disease states, rather than treatment conditions. These data suggest a longer-term, intrinsic conservation of the T2D circadian signature in primary myocytes. This finding would appear consistent with data demonstrating that induced pluripotent stem cells from donors with T2D that were differentiated into myoblasts had multiple defects as compared to healthy subjects, including reduced insulin-stimulated glucose uptake and reduced mitochondrial oxidation (Batista et al., 2020). These T2D-associated defects were conserved despite the robust manipulations that the cells undergo. How cultured myocytes from T2D donors preserve a dysfunctional phenotype, including insulin resistance (Bouzakri and Zierath, 2007; Sarabia et al., 1992), has not been fully elucidated, but likely reflects genetic background and epigenetic mechanisms. Our data reveal that an intrinsic disruption of circadian biology in skeletal muscle may predispose to insulin resistance and metabolic disease. The cause of this disruption is unknown, but in addition to inherited factors, cumulative effects of sedentary lifestyle, sleep deprivation, nutritional and hormonal factors are likely to play a role (Gabriel and Zierath, 2019; Zimmet et al., 2016).

Our results demonstrate that dysregulated rhythmic mitochondrial metabolism may play a role in mediating the disrupted rhythmic cellular metabolism and molecular-clock machinery in primary myocytes from individuals with T2D. Mitochondrial diurnal rhythms in oxidative capacity and oxygen metabolism have been demonstrated in several tissues, including skeletal muscle *in vivo* (Hansen et al., 2016). These skeletal muscle diurnal mitochondrial rhythms are more likely to emanate from mitochondrial membrane dynamic processes, regulated by rhythmic proteins such as OPA1 and FIS1, rather than a day-night rhythm of mitochondrial biogenesis and content (de Goede et al., 2018; van Moorsel et al., 2016). Mitochondrial membrane dynamics demonstrate circadian behavior in several peripheral tissues (de Goede et al., 2018), however, transcriptional control of these pathways is tissue-specific (Schrauwen et al., 2010; Weaver et al., 2014). For example, while *Drp-1*, *Mfn1/2*, and *Opa1* are not direct targets of the clock machinery in liver (Jacobi et al., 2015), *Mfn1* and *Opa1* are decreased in cardiac muscle of *Bmal1^-/-^* mice (Kohsaka et al., 2014).

In the current study, ChIP-sequencing revealed that the inner-mitochondrial genes highly associated with insulin sensitivity also were direct targets of both CLOCK and BMAL1 in skeletal muscle. Moreover, some of these genes were also circadian in myotube cultures from donors with T2D. However, genes with CLOCK and BMAL1 binding did not coalesce with inner-mitochondrial genes in myotubes from donors with NGT, despite the similar enrichment of these pathways when assessing genes with reduced rhythmic amplitudes in T2D and genes that correlated with insulin sensitivity *in vivo.* Mitochondrial metabolism plays a role in bi- directional regulation of diurnal metabolism in peripheral tissues (Manella and Asher, 2016). Here we report that myotube cultures from donors with NGT displayed more robust mitochondrial metabolic rhythms as compared with T2D. Mitochondrial rhythms in individuals with NGT may help to maintain the diurnal metabolic rhythm in skeletal muscle. Additionally, manipulating mitochondrial membrane metabolism may alter the rhythmic expression of molecular-clock genes (Lassiter et al., 2018; Schmitt et al., 2018), although the signaling mechanism between mitochondrial metabolism and the molecular-clock has not been elucidated. Our results provide evidence that *OPA1* depletion disrupts molecular-clock genes *NPAS2, PER2* and *PER3* in primary myocytes.

Mitochondrial respiration and ROS production are tightly linked, and alongside mitochondrial capacity and oxygen consumption, antioxidant proteins and ROS production also display circadian activity (de Goede et al., 2018). In our study, mRNA expression of antioxidant enzymes that localize to mitochondria (*GPX1, GPX3, GPX4*) were rhythmic in myotube cultures from donors with NGT, but not T2D. The aforementioned enzymes are all involved in the detoxification of hydrogen peroxide, a known regulator of HIF1-α activity (Hoppeler et al., 2003), which was also rhythmic in myotube cultures from donors with NGT, but not T2D. HIF1-α plays a key role in mediating signaling from the core-clock to mitochondria and vice-versa in skeletal muscle. Our data indicate that altered rhythms of mitochondrial respiration and ROS handling in myotube cultures from donors with T2D may be partly responsible for the ablated rhythm of HIF1-α expression. In support of this, the OPA1-mediated increase in *NPAS2* expression and mitochondrial ROS were rescued by treatment with resveratrol, an antioxidant and HIF1-α modulator. Collectively, our data show cross-talk between the inner-mitochondrion and the molecular-clock, with genes of these pathways rhythmically dysregulated in T2D. Dysfunction of the inner-mitochondrion in peripheral tissues is implicated in the pathogenesis of insulin resistance and T2D, concomitant with increased ROS production, decreased oxidative capacity, and reduced metabolic flexibility (Schrauwen et al., 2010). This dysfunction may also play a role in regulating intrinsically altered metabolic, and molecular-clock rhythms in T2D.

In conclusion, we show disturbances in the intrinsic rhythmicity of gene expression and metabolism in skeletal muscle cells of individuals with T2D. This dysregulation is evident in myocytes in the absence of systemic factors, hormones, nutritional cues, or direct influence of the central or other peripheral clocks. Moreover, circadian inner-mitochondrion gene amplitudes are downregulated in T2D. These genes are also associated with insulin sensitivity and are under bi-directional regulation of circadian metabolism in skeletal muscle. The dysregulation of circadian metabolism in skeletal muscle of people with T2D underscores the need to take circadian biology into account and consider approaches in chrono-medicine (Roenneberg and Merrow, 2016) when prescribing pharmacological therapy, particularly treatments that affect mitochondrial function (Fontaine, 2018). Our findings provide mechanistic insight into T2D pathophysiology and have clinical implications into the link between insulin sensitivity and environmental triggers that are associated with altered metabolism, including impaired sleeping-patterns, social jet-lag, or shift-work.

## Acknowledgments

The authors are supported by grants from the AstraZeneca SciLifeLab Research Programme, Novo Nordisk Foundation (NNF14OC0011493, NNF14OC0009941, NNF17OC0030088), Swedish Diabetes Foundation (DIA2018-357), Swedish Research Council (2015-00165, 2018- 02389), the Strategic Research Programme in Diabetes at Karolinska Institutet (2009-1068), the Stockholm County Council (SLL20170159) and the Swedish Research Council for Sport Science (P2019-0140). Brendan M. Gabriel was supported by fellowships from the Novo Nordisk Foundation (NNF19OC0055072), the Wenner-Gren Foundation, and an Albert Renold Travel Fellowship from the European Foundation for the Study of Diabetes and an Eric Reid Fund for Methodology from the Biochemical Society. Nicolas J. Pillon and Laura Sardon Puig were supported by an Individual Fellowship from the Marie Skłodowska-Curie Actions (European Commission: 704978, 675610). Xiping Zhang and Karyn Esser were supported by NIH R01AR066082. Nicolas J. Pillon were supported by grants from the Sigurd och Elsa Goljes Minne and Lars Hierta Memorial Foundations (Sweden). We acknowledge the Beta Cell in-vivo Imaging/ Extracellular Flux Analysis core facility supported by the Strategic Research Program in Diabetes for the usage of the Seahorse flux analyzer. Additional support was received from the Novo Nordisk Foundation Center for Basic Metabolic Research at the University of Copenhagen (NNF18CC0034900). The Novo Nordisk Foundation Center for Basic Metabolic Research is an independent research center at the University of Copenhagen, partially funded by an unrestricted donation from the Novo Nordisk Foundation. We acknowledge the Single-Cell Omics platform at the Novo Nordisk Foundation Center for Basic Metabolic Research for technical and computational expertise and support.

## Author Contributions

Conceptualization: BMG, AA, MR, KAE, AK, JRZ

Methodology: BMG, AA, LSP, JABS, NJP, ZL, JTL AZ, KAE, RB, MR, AK, JRZ

Formal Analysis: BMG, AA

Investigation: BMG, AA, LSP, JABS, ZL, LD, AA, XZ, ALB, RCL, NJP Resources: JTT, AZ, JTL, KAE, RB, MR, NJP, AK, JRZ

Writing – Original Draft: BMG, JRZ

Writing –Review & Editing: BMG, AA, LSP, JABS, XZ, ALB, RCL, HG, ZL, LD, JTT, AZ, ZH, MR, JTL, KAE, RB, NJP, AK, JRZ

Funding Acquisition: BMG, NJP, JTT, JTL, AZ, KAE, RB, MR, AK, JRZ

## KEY RESOURCES TABLE

| REAGENT or RESOURCE | SOURCE | IDENTIFIER |
| --- | --- | --- |
| **Antibodies** |  |  |
| Anti-OPA1 | Cell Signaling Technology (USA) | #80471 |
| Total OXPHOS | Abcam (UK) | #110411 |
| β-actin | Sigma-Aldrich (USA) | #A5441 |
| BMAL1 | Sigma-Aldrich (USA) | #SAB4300614 |
| CLOCK | Abcam (USA) | #ab3517 |
| **Biological Samples** |  |  |
| Human skeletal muscle biopsies | Karolinska Institutet | Described in methods |
| Myo-Cre Opa1<sup>-/-</sup> mice | University of Barcelona | Described in methods |
| C57Bl/6j mice skeletal muscle | University of Florida | Described in methods |
| **Critical Commercial Assays** |  |  |
| Seahorse XF Cell Mito Stress Test Kit | Agilent Technologies, USA | Kit 103015-100 |
| **Deposited Data** |  |  |
| Raw and analyzed data | This paper | GEO<br>https://www.ncbi.nlm.nih.gov/geo/query/acc.cgi?acc=GSE143334, |
| **Experimental Models: Cell Lines** |  |  |
| Primary human skeletal muscle cells normal glucose tolerant | Karolinska Institutet | Described in methods |
| Primary human skeletal muscle cells Type 2 diabetes | Karolinska Institutet | Described in methods |
| **Oligonucleotides** |  |  |
| Primers for RT-qPCR: see table 3 of methods | This paper | Described in Table 3 of methods |
| **Software and Algorithms** |  |  |
| Prism: version 8 | Graphpad software | N.A. |
| R: version 3.5 | N.A. | N.A. |
| Python version 3.7 | N.A. | N.A. |
| Microsoft Excel: version 2016 | Microsoft | N.A. |
| FastQC (version 0.11.5) | Babraham Institute, UK | link |
| Trim_Galore! (version 0.4.3) | Babraham Institute, UK | link |
| STAR (version 2.5.3a) | Dobin et al., 2013 | PMID: 23104886<br>DOI:<br>[10.1093/bioinformatics/bts635](https://doi.org/10.1093/bioinformatics/bts635) |
| featureCounts (version 1.5.2) | Liao et al., 2014 | PMID: 24227677<br>DOI:<br>10.1093/bioinformatics/btt656 |
| multiQC (version 1.5) | Ewls et al., 2016 | PMID: 27312411<br>DOI:<br>10.1093/bioinformatics/btw354 |
| limma (version 3.42.2) | Ritchie et al., 2015 | PMID: 25605792<br>DOI:<br>10.1093/nar/gkv007 |
| DODR (version 0.99.3) | Thaben &<br>Westermarck, 2016 | PMID: 27207944,<br>DOI: 10.1093/bioinformatics/btw309 |
| RAIN (version 1.20.0) | Thaben &<br>Westermarck, 2014 | PMID: 25326247,<br>DOI:<br>10.1177/0748730414553029 |
| HarmonicRegression (version 1.0) | Lück et al., 2014 | PMID: 25373909<br>DOI:<br>10.1016/j.celrep.2014.09.021 |
| clusterProfiler (version 3.14.3) | Yu et al. 2012 | PMID: 22455463<br>DOI:<br>10.1089/omi.2011.0118 |
| Other |  |  |

## Methods

### Resource Availability Lead Contact

Further information and requests for resources and reagents should be directed to and will be facilitated by the Lead Contact, Juleen R. Zierath.

### Materials Availability

This study did not generate new unique reagents.

### Data and Code Availability

The data and codes related to the findings of this study will be available from the corresponding author after publication upon reasonable request. The research team will provide an email address for communication once the data are approved to be shared with others. The proposal with detailed aims, statistical plan, and other information/materials may be required to guarantee the rationality of requirement and the security of the data. The patient-level data, but without names and other identifiers, will be shared after review and approval of the submitted proposal and any related requested materials.

### Human study methods

#### Study group and skeletal muscle biopsy procedure

Study groups included individuals with normal glucose tolerance (NGT), as determined by oral glucose tolerance test (OGTT), or Type 2 diabetes (T2D). Written informed consent was obtained from all participants. The study protocol was approved by the regional ethics board in Stockholm, Sweden. The participants were instructed to refrain from physical exercise 48 hrs before the visit. All investigations were performed in the morning after an overnight fast. Anthropometric measurements were taken, and blood samples were obtained for clinical chemistry analysis. After application of local anesthesia (10 mg/ml mepivacaine hydrochloride, AstraZeneca, Cambridge, UK), a biopsy was obtained from m. *vastus lateralis* using a Weil-Blakesley conchotome instrument (Agnthos, Sweden). Biopsies were cleared from any visible fat, connective tissue, or blood vessels and either used directly to prepare primary muscle cell cultures or immediately frozen in liquid nitrogen and stored at −80°C until microarray analysis. The clinical characteristics of the NGT (*n*=7) or T2D (*n*=5) donors for the muscle cell culture studies are presented in Table 1.

**Table 1.** Clinical Characteristics of the Donors for Muscle Cell Culture Studies.

|  | NGT |  | T2D |  | p-value |
| --- | --- | --- | --- | --- | --- |
|  | Mean | SEM | Mean | SEM |  |
| **Age** | 62.6 | 1.1 | 60 | 1.4 | **0.005** |
| **Weight [kg]** | 93.7 | 3.6 | 89.7 | 6.3 | 0.174 |
| **Height [cm]** | 180.3 | 1.9 | 180.0 | 3.6 | 0.875 |
| **BMI [kg/m<sup>2</sup>]** | 28.8 | 0.8 | 27.6 | 1.2 | **0.012** |
| **BP Systolic [mm Hg]** | 139.6 | 3.3 | 138.2 | 8.9 | 0.716 |
| **BP Diastolic [mm Hg]** | 87.9 | 3.0 | 81.5 | 5.4 | **0.022** |
| **Pulse [BPM]** | 61.5 | 4.5 | 65.3 | 4.1 | 0.099 |
| **HbA1c [mmol/mol]** | 30.3 | 1.5 | 59.6 | 5.6 | **<0.0001** |
| **Plasma-glucose 0 OGTT [mmol/l]** | 5.14 | 0.20 | 9.06 | 0.85 | **<0.0001** |
| **Plasma-glucose 120 OGTT [mmol/l]** | 5.96 | 0.67 | 16.16 | 1.39 | **<0.0001** |
| **Plasma-TG [mmol/l]** | 1.41 | 0.38 | 1.25 | 0.26 | 0.084 |
| **Plasma-Chol [mmol/l]** | 5.83 | 0.37 | 3.84 | 0.36 | **<0.0001** |
| **Plasma-HDL [mmol/l]** | 1.37 | 0.16 | 1.22 | 0.08 | **<0.0001** |
| **Plasma-LDL [mmol/l]** | 3.83 | 0.35 | 2.00 | 0.25 | **<0.0001** |
| **VO<sub>2</sub>max [ml/min]** | 2478 | 207 | 2735 | 361 | 0.194 |
| **VO<sub>2</sub>max [ml/kg/min]** | 28.7 | 2.0 | 31.6 | 2.6 | 0.153 |
| **Max workload [Watt]** | 181.9 | 16.8 | 193.8 | 26.3 | 0.454 |

Skeletal muscle biopsies were obtained from Normal Glucose Tolerant (NGT; n=7) and Type 2 Diabetic (T2D; n=5) men and myotube cultures were derived for *in vitro* studies. Results are mean±SEM. P-values were determined by Student's *t*-test.

### Cell culture

Primary myoblasts from muscle biopsies from men with NGT and T2D were grown in DMEM/F12+Glutamax with 16% FBS and 1% Antibiotic-Antimycotic (100X). Differentiation was induced at 80% confluence, by incubating the cells for 4-5 days in fusion medium consisting of 76% DMEM Glutamax with 25 mmol/l glucose, 20% M199 (5.5 mmol/l), 2% HEPES and 1% Antibiotic-Antimycotic (100X), with 0.03 μg/ml zinc sulfate and 1.4 mg/ml Vitamin B12, with a further addition of 100 μg/ml apo-transferrin and 0.286 IU/ml insulin prior to use. Experiments were performed on differentiated myotubes after an additional 3-5 days incubation in the same media containing 2% FBS (post fusion media).

### Circadian experiments

After post fusion, DMEM with 5.5 mmol/l glucose was used in the post fusion medium to create a low glucose condition. After 22 hrs, cells were synchronized by serum shock (media containing 50% FBS, 2 hrs) (Balsalobre et al., 1998) and then switched back to low glucose media. Skeletal muscle myotubes from men with either T2D or NGT were lysed at 6-hr intervals from 12 hrs to 54 hrs. Zeit-time (ZT) was determined as time after synchronization.

### RNA-sequencing

RNA was checked for quality using the Agilent RNA 600 nano kit and Bioanalyser instrument (Agilent Technologies, Santa Clara, CA). Aliquots of RNA (1 µg) were analyzed using the Illumina TruSeq Stranded Total RNA with Ribo-Zero Gold protocol (Illumina) as described (Laker et al., 2017; Nylander et al., 2016). Ribosomal RNA was removed from the sample using 35 µl rRNA removal beads (Illumina) on a magnetic plate followed by clean-up of the ribosomal-depleted RNA with 193 µl Agencourt RNAClean XP beads (Beckman Coulter), 70% ethanol wash and elution into 10 µl Elution buffer (Illumina). The RNA sample was fragmented for 4 min at 94°C in Elute, Prime, Fragment High Mix (Illumina) and then subjected to first strand cDNA synthesis with 1 µl Superscript III reverse transcriptase (Life Technologies) per sample using a thermocycler programmed to 25°C for 10 min, 50°C for 15 min and 70°C for 15 min. Second strand cDNA was synthesized by addition of Second Strand Marking Master Mix and samples were incubated at 16°C for 60 min. Samples were subject to another bead clean up prior to A-tailing and ligation of adapters as per kit instructions (Illumina). Following a third bead clean-up, samples were enriched for DNA fragments by amplification using the Illumina PCR Primer Cocktail and PCR Master Mix, subjected to 98°C for 30 min, followed by a pre-defined cycle (98°C for 10 s, 60°C for 30 s and 72°C for 30 s) that was repeated 3-15 times, based on each individual sample, and finally incubated for 5 min at 72°C. Samples were cleaned, and validated for DNA concentration using the Qubit dsDNA HS assay kit (Invitrogen) and for base pair size and purity using the Agilent High Sensitivity DNA chip and Bioanalyser instrument. Libraries were subjected to 100-bp single-end sequencing on the X Ten platform (Illumina) at the Beijing Genomics Institute (BGI), Hong Kong, China. Approximately 20 million reads/sample were assigned to genes with 14,051 genes surviving the expression threshold.

### RNA-seq data analysis

RNA-seq reads (*n̄* ≈ 40 M) from FASTQ files were quality-trimmed using Trim_Galore (v0.4.3). Trimmed reads were aligned using STAR (v2.5.3a) (Dobin et al., 2013)aligner with Ensembl human annotation (GRCh38, release 92) (Zerbino et al., 2018) and gene features were counted using featureCounts from subread (v1.5.2) package (Liao et al., 2014) resulting in 27 M uniquely mapped and 20 M assigned reads to genomic features (genes) on average, respectively. The lowly expressed genes were discarded from downstream analysis using filterByExpr function from edgeR package (Robinson et al., 2010) resulting 18,482 genes. As an apparent batch effect was introduced by participants (Figure S1), this batch effect was removed by using limma’s removeBatchEffect function (Ritchie et al., 2015) for further downstream rhythmicity (RAIN) and differential rhythmicity (DODR) analysis as these tools do not consider any batch effect by default. The batch corrected CPM values were in log2 scale.

### Rhythmicity and differential rhythmicity analysis

*p* values for rhythmicity were assessed using RAIN (Thaben and Westermark, 2014) with longitudinal method and adjusted for multiple testing using Benjamini-Hochberg method. Genes with adjusted p-value below 0.10 (FDRRAIN < 0.10) were considered as rhythmic. We chose RAIN because this algorithm detects both symmetric and non-symmetric wave forms (Thaben and Westermark, 2014). Differential rhythmicity analysis was performed on mean centered logCPM values (batch corrected) between myotubes from NGT and T2D donors (control and HGI group, separately) by using DODR (Thaben and Westermark, 2016) with “robust” method and resulting meta p-values were adjusted for multiple testing using Benjamini-Hochberg. It is essential to mean-center the logCPM values as since DODR internally assumes identical means in order to test for differential rhythmicity. Genes with adjusted meta p-value below 0.10 (FDRDODR < 0.10) were considered differentially rhythmic.

### Circadian amplitude estimations

A harmonic regression model using HarmonicRegression R package (Luck et al., 2014) with first degree polynomial component was fit to (participant) batch corrected logCPM values. As the calculated absolute amplitudes were in log2 scale (*Alog*), these amplitudes were transformed to relative amplitudes using the following equations. As fold-change amplitude (*Afc*) is

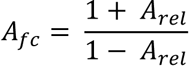

and twice the absolute amplitude calculated from log2 transformed data (batch corrected logCPM) equals to log2 of fold-change amplitude (*Afc*):

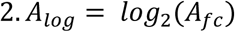

the relative amplitude (*Arel*) can be derived by solving these two equations to explain the relationship between *Alog* and *Arel* as

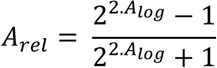

To compare the relative amplitudes between T2Ds and NGTs, log2 ratios were calculated for each treatment group (control and HGI):

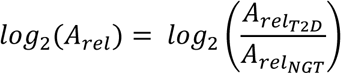

### Gene enrichment analysis

Gene enrichment analysis was performed by using both over-representation analysis (ORA) and gene-set enrichment analysis (GSEA) throughout this study. For biological interpretation of the biological processes, we used Reactome pathways (Jassal et al., 2020) while gene ontology (GO) (Ashburner et al., 2000) cellular components (CC) was used in order to identify the enrichment of subcellular compartments. clusterProfiler (Yu et al., 2012) R package was used to analyze all gene enrichment results. Ontology terms (Reactome, GO:CC) with an adjusted p-value, using Benjamini-Hochberg method, below 0.10 were considered to be significantly enriched.

As we aimed to investigate the important biological pathways associated with the circadian genes, we subsetted circadian genes by disease-treatment groups (Figure 1D) and peak times (Figure 2F) and analyzed enrichment from Reactome pathways using ORA. Similarly, genes with significant correlations between insulin resistance (M-value) and basal gene expression (Figure 4C) were tested for GO:CC enrichment using ORA method. When a ranking metric, such as relative amplitude log2-fold-changes (Figure 4C), was preferred to analyze enrichment for cellular components, we used GSEA method with GO:CC database. An ontology-gene interaction network with top enriched (GSEA) GO:CC terms, so called CNET plot, was generated to show the involvement of an individual gene, together with the Spearman correlation coefficient (Figure 4D) and relative amplitude log2-fold-change (Figure 3B).

Lastly, clustering analysis revealed 10 distinct clusters from integrative analysis using binary input for presence/absence of rhythmicity in disease-treatment groups, insulin sensitivity correlation with basal gene expression and BMAL/CLOCK binding. The genes in each cluster were analyzed using GO:CC database with ORA method (Figure 5C), similar to the analysis performed in Figure 1D and 2F.

#### Hyperinsulinemic euglycemic clamp procedure

This study group was handled as described in the section “Human study methods”. The clinical characteristics of the men with NGT (*n*=22) or T2D (*n*=22) enrolled in the gene array and hyperinsulinemic euglycemic clamp study are reported in Table 2. After a 45 min rest, the study participants underwent a hyperinsulinemic euglycemic clamp as described (Andersson et al., 2014). Following an intravenous bolus dose of insulin (1,600 mU/m^2^ body surface area), insulin was infused intravenously at a rate of 40 mU/m^2^/min for 2 hrs and a variable intravenous infusion of glucose (200 mg/ml) was used to maintain euglycemia between 81-99 mg/dl (4.5-5.5 mmol/l). The infusion rate of glucose during the last 60 min of the clamp, when insulin levels are in a steady state, was used to calculate whole-body glucose disposal rates (M-value).

**Table 2.** Clinical Characteristics of the Participants in the Hyperinsulinemic Euglycemic Clamp Study.

|  | NGT |  | T2D |  | p-Value |
| --- | --- | --- | --- | --- | --- |
|  | Mean | SEM | Mean | SEM |  |
| Age | 57.6 | 2.2 | 61.8 | 1.5 | 0.116 |
| Weight [kg] | 83.5 | 2.2 | 88.0 | 1.7 | 0.116 |
| Height [m] | 1.8 | 0.0 | 1.8 | 0.0 | 0.090 |
| BMI [kg/m<sup>2</sup>] | 26.8 | 0.4 | 27.3 | 0.5 | 0.398 |
| Waist [cm] | 95.9 | 1.9 | 98.7 | 1.3 | 0.239 |
| Hip [cm] | 100.1 | 1.1 | 102.0 | 1.1 | 0.238 |
| W/H ratio | 1.0 | 0.0 | 1.0 | 0.0 | 0.409 |
| Body fat [%] | 22.8 | 1.0 | 24.0 | 1.0 | 0.399 |
| BP Systolic [mm Hg] | 138.2 | 2.4 | 143.6 | 2.8 | 0.140 |
| BP Diastolic [mm Hg] | 83.3 | 1.3 | 83.9 | 1.7 | 0.770 |
| Pulse [BPM] | 62.7 | 2.2 | 67.3 | 1.5 | 0.089 |
| Fasting plasma-Glucose [mmol/l] | 5.4 | 0.1 | 7.8 | 0.3 | **0.000** |
| HbA1c [mmol/mol] | 36.6 | 0.7 | 50.9 | 1.2 | **0.000** |
| Fasting serum-Insulin [pmol/l] | 48.7 | 6.6 | 71.3 | 7.7 | **0.030** |
| HOMA1-IR | 1.8 | 0.3 | 3.6 | 0.4 | **0.001** |
| M-Value [μmol/(kg*min)] | 34.2 | 3.1 | 15.3 | 1.9 | **0.000** |
| Creatinine [μmol/l] | 85.2 | 1.8 | 82.6 | 3.2 | 0.490 |
| ASAT [μkat/l] | 0.4 | 0.0 | 0.4 | 0.0 | 0.694 |
| ALAT [μkat/l] | 0.4 | 0.0 | 0.5 | 0.0 | 0.407 |
| Plasma TG [mmol/l] | 1.1 | 0.2 | 1.2 | 0.2 | 0.676 |
| Plasma Chol [mmol/l] | 5.3 | 0.2 | 4.5 | 0.2 | **0.005** |
| HDL Chol [mmol/l] | 1.4 | 0.1 | 1.3 | 0.1 | 0.137 |
| LDL Chol [mmol/l] | 3.6 | 0.2 | 2.6 | 0.2 | **0.001** |
| FFA [mmol/l] | 0.5 | 0.0 | 0.5 | 0.0 | 0.433 |
| C-Peptide [nmol/l] | 0.7 | 0.0 | 0.9 | 0.1 | **0.007** |

The study groups include men with normal glucose tolerance (NGT; n=25), as determined by oral glucose tolerance test, or T2D (n=25). Skeletal muscle biopsies were obtained for gene expression analysis. Insulin sensitivity (M-value) was determined by the hyperinsulinemic euglycemic clamp. Results are mean ± SEM. P-values were determined by Student's *t*-test.

### ChIP-sequencing

#### Tissue harvesting

Animal procedures were conducted in accordance with institutional guidelines for the care and use of laboratory animals as approved by the University of Florida Institutional Animal Care and Use Committee. For the ChIPseq data, two replicate samples for BMAL1 and CLOCK were used. Each sample required pooling gastrocnemius muscle from 10 adult C57BL/6J male mice (Jackson Labs, Farmington, Connecticut, USA). The mice were entrained to 12L/12D schedule and all tissues were collected at ZT2 and frozen immediately.

#### ChIP-Seq sample preparation

For the CLOCK ChIPseq samples: Skeletal muscle was homogenized (muscle:buffer ratio 1:10) in lysis buffer (10 mM HEPES, pH7.5, 10 mM MgCl2, 60 mM KCl, 300 mM sucrose, 0.1 mM EDTA, pH 8.0, 0.1% Triton X-100, 1 mM DTT) with EGS (ethylene glycol bis (succininic acid) N-hydroxysuccinimide ester, Bio-world) (Koike et al., 2012). The reaction was stopped with 1 M Tris (pH 7.5) buffer (final concentration of Tris was 20 mM) and formaldehyde added (final concentration of 1%). Samples were incubated for 25 min at room temperature and crosslinking was stopped by adding glycine to 125 mM.

For the BMAL1 ChIPseq samples, we followed the protocol outlined in Hodge et al., 2019 (Hodge et al., 2019). Skeletal muscles were homogenized in lysis buffer containing 1% of formaldehyde. Samples were incubated at room temperature for 25 min and crosslinking was stopped by adding glycine to 125 mM.

For all samples, myonuclei were isolated from skeletal muscle homogenates as described (Hodge et al., 2019) and samples were incubated with the appropriate antibody at 4°C overnight. Samples were washed and then incubated at 65°C overnight to de-crosslink. Lastly, RNase, 5 M NaCl and Proteinase K were added, and samples were incubated at 55°C for 1 hr. DNA was recovered with a PCR purification kit (Qiagen) and the ChIPseq library was prepared by using NEBNext Ultra DNA library prep kit for Illumina. Library quality was determined by Tapestation and Quantitative PCR. Sequencing parameters were paired end 100 bp run with approximately 40M reads/sample on the HiSeq 3000 Illumina sequencing system.

#### ChIP-Seq data analysis

The reads were aligned to *Mus musculus* genome assembly GRCm38 (mm 10) using Bowtie2 (Langmead and Salzberg, 2012) with the --sensitive-local option, which does not require that the entire read aligns from one end to the other. The biological replicates of the aligned reads were merged for *CLOCK*, *BMAL1*, and input, respectively. Homer software (Heinz et al., 2010) was deployed to perform peak calling for the CLOCK sample and the BMAL1 sample with the input sample as background. The default FDR rate threshold 0.001 was used to detect significant peaks.

### Integrative analysis of circadian rhythmicity, insulin sensitivity and BMAL1/CLOCK chromatin binding

In order to identify the gene patterns across different experimental procedures, an integrative analysis approach was applied to circadian genes (RNAseq), insulin sensitivity correlations (M-value vs. basal gene expression from microarray study) and BMAL1/CLOCK chromatin binding (ChIP-seq). Both gene expression correlations with M-values and BMAL1/CLOCK ChIP-seq experiments were performed on mice, therefore we have identified the homologous genes between mouse and human and only included those genes for further downstream analysis. ChIP-seq peaks closest to a gene’s transcription start site (TSS) are assigned to that gene. A binary matrix was created with the circadian rhythmicity (1: rhythmic, 0: non-rhythmic), Spearman correlation between insulin sensitivity (M-value) and basal gene expression (1: significant, 0: non-significant) and BMAL1/CLOCK binding (1: bound, 0: not-bound). The resulting matrix was used for clustering analysis using “clust” algorithm (Abu-Jamous and Kelly, 2018) in order to identify groups of genes showing similar patterns. This resulted in 10 different clusters. Those 10 clusters were used for gene enrichment analysis using ORA method and GO:CC database.

The circadian genes associated with BMAL1/CLOCK binding were identified by overlapping each group, NGTs and T2Ds with both treatment conditions separately. The overlaps were tested by Fisher’s exact test and resulting p-values were adjusted by Benjamini-Hochberg method. Adjusted p-values below 0.05 were considered as statistically significant. Overlapping gene percentage, also known as Jaccard index, was used for showing the similarity between the overlaps (Figure 5A).

### Animal models

#### Myo-Cre Opa1^-/-^ mice

Animal work was performed in compliance with guidelines established by the University of Barcelona Committee on Animal Care. *Opa1^loxP/loxP^* mice were generated as reported (Ramirez et al., 2017; Rodriguez-Nuevo et al., 2018). The *Myo-Cre OPA1* mouse line was generated by crossing homozygous *OPA1^loxP/loxP^* mice with a strain expressing *Cre* recombinase under the control of the myogenin promoter and the 1-kb mouse MEF2C enhancer, thus yielding a transgene called *Myo*-*Cr*e (Li et al., 2005). *Cre OPA1* littermates were used as controls for knockout (KO) mice. Mice were kept under a 12-hr dark/light cycle and were provided standard chow diet and water *ad libitum*. Four-month-old male mice were used in all experiments. On the experimental day, mice were anesthetized with isoflurane, killed by cervical dislocation, and *tibialis anterior* muscles were harvested, and immediately frozen in liquid nitrogen for subsequent analysis.

### Mitochondrial disrupting compounds and circadian studies

Primary human skeletal muscle cells from men with NGT were used for the study of the effect of compounds FCCP, Rotenone a, antimycin a, and oligomycin on molecular clock gene expression. Primary skeletal muscle cells were grown and differentiated as described above. A Lactate Dehydrogenase (LDH) assay was performed to determine cell viability after treatment with compounds, there were no differences in LDH concentration in any condition (data not shown). The cells were synchronized as described above and compounds or vehicle control were added at ZT14 after synchronization, and cells harvested after 4h exposure (ZT18).

Compounds were Rotenone/Antimycin A (0.38 µm), FCCP (2 µm) and Oligomycin (1 µm). Cells were harvested to collect RNA and core-clock gene mRNA expression was measured by RT-qPCT.

### Gene silencing and metabolic studies

#### Opa1 silencing

Primary human skeletal muscle cells from men with NGT were used for the study of *OPA1* silencing. Primary skeletal muscle cells were grown and differentiated as described above. After switching to post fusion media, cells were transfected as described (Lassiter et al., 2018), using 5 µM of either Silencer® Select (Thermo Fisher Scientific, Waltham, MA) Negative Control No. 2 (#4390847) or validated siRNAs against *OPA1* (#s9852), for two separate 5-hr transfection periods separated by ∼48 hrs.

#### Metabolic phenotyping of cells

To assess the mitochondrial function of primary human skeletal muscle cells, cells were subjected to a Seahorse XF Mito Stress Test using the manufacturer’s instructions for timing (Agilent, Santa Clara, CA), as described (Lassiter et al., 2018). Oxygen consumption rates (OCR) and extracellular acidification rates (ECAR) were measured at three time points under unstimulated conditions, then after treatment with 1 µM oligomycin, 2 µM FCCP, and 0.75 µM rotenone + antimycin A. OCR and ECAR was normalized to the protein concentration, as quantified by a Pierce BCA protein assay kit (Thermo Fisher Scientific). Myotube cultures either from men with NGT (*n*=5) or T2D (*n*=5) were subjected to a Mito Stress (Figure S1) test under the same conditions as previously described, except, these cells were measured at ZT24 after synchronization as described in ‘Circadian experiments’ section. For circadian Seahorse XF experiments (time-course of basal OCR, Figure 3D), myotube cultures either from men with NGT (*n*=5) or T2D (*n*=5), were synchronized as described in “Circadian experiments” section. Myotubes were measured sequentially, i.e. the same cells were measured at all time points in buffered media, and only basal oxygen consumption was measured. This allowed minimal disturbance to cells, and enabled measurements of diurnal oxygen consumption in a method derived from previous work (Peek et al., 2013).

#### NAD measurements of cells after OPA1 silencing

NAD levels were assessed in primary human skeletal muscle cells after *OPA1* silencing using an enzymatic cycling assay (Dall et al., 2018). Cells were harvested with trypsin, and the cell pellet was dissolved in 200 µl of 0.6 M perchloric acid and subjected to centrifugation for 3 mins at 13,000g. The supernatants were transferred to new tubes and diluted 1:100 in 100 mM Na2HPO4 (pH 8). An aliquot (100 µl) of the diluted extract was pipetted into a 96-well plate, and a reaction mix (100 µl) containing 100 mM Na2HPO4, 10 µM flavinmononucleotide, 2% ethanol, 90 U/ml alcohol dehydrogenase, 130 mU/ml diaphorase, 2.5 µg/ml resazurin and 10 mM nicotinamide was added. Fluorescence (Ex 540 nm/ Em 580) was measured over 30 min and the NAD content was calculated from a standard curve and normalized to the protein concentration. Protein was extracted by dissolving the cell pellet (after perchloric acid precipitation) in 200 µl 0.2M NaOH, incubating at 95°C for 5 min, and centrifugation for 5 mins at 13,000 *g*. Protein concentration was determined on the supernatant using the Bicinchoninic Acid Assay (23227, Thermo Fisher Scientific, MA).

### Resveratrol treatment

Human primary skeletal muscle cells were grown, differentiated and synchronized as described above. Cells were either treated with 10 µM resveratrol (#R5010, Merck KGaA, Darmstadt, Germany) or vehicle control for the duration of the experiment, subsequent to synchronization.

### Gene expression analysis

#### Microarray analysis

RNA quality was assessed and ensured using the Experion™ Automated Electrophoresis System (Bio-Rad Laboratories, CA). Affymetrix GeneChip™ Human Transcriptome Array 2.0

(Thermo Fisher Scientific, MA) was used for the whole-transcriptome analysis performed at the Bioinformatics and Expression Core facility (BEA) at Karolinska Institutet, Huddinge, Sweden. Amplified and biotinylated sense-strand DNA targets were generated from total RNA using the WT Plus Kit (Thermo Fisher, MA). Fragmented and biotinylated sense-strand DNA target (5.5 µg) were hybridized to the arrays in GeneChip Hybridization Oven 645 at 45°C for 16-18 hrs. Arrays were washed and stained in GeneChip Fluidics Station 450 prior to scanning in Affymetrix GeneChip Scanner. Gene array data were analyzed with packages available from Bioconductor (http://www.bioconductor.org). Normalization and calculation of gene expression was performed with the Robust Multichip Average expression measure using oligo package (Carvalho and Irizarry, 2010). Prior to further analysis, a nonspecific filter was applied to include genes with expression signal >30 in at least 25% of all samples Differentially expressed genes between NGT and T2D were identified using limma (Ritchie et al., 2015). The correlation between the basal gene expression and insulin sensitivity (M-value) was calculated by Spearman’s rank correlation and p-values are adjusted for multiple testing using Benjamini-Hochberg method. Spearman correlation coefficients were used for gene enrichment analysis using GSEA method with GO:CC database (See section “Gene enrichment analysis” for details). Ontology-gene interaction network, so called CNET plot, from this analysis using the top enriched terms are shown in Figure 4D. The color code represents the log2-fold-change of gene expression between NGT compared to T2D.

#### PCR-analysis

RNA was prepared from human skeletal muscle biopsies and primary human skeletal muscle cells. Gene expression was determined by using Fast SYBR Green Master Mix (Thermo Fisher Scientific) and self-designed oligonucleotides (oligonucleotide sequences provided below) or Predesigned TaqMan Gene Expression Assays (Thermo Fisher Scientific). SYBR human housekeeping: *B2M*, *GUSB*, *TBP*, *RPLO*. TaqMan mouse housekeeping genes: *B2m* and *Gapdh*, human: *TBP, B2M.* Primer sequences and assay IDs presented in Table 3.

**Table 3.**
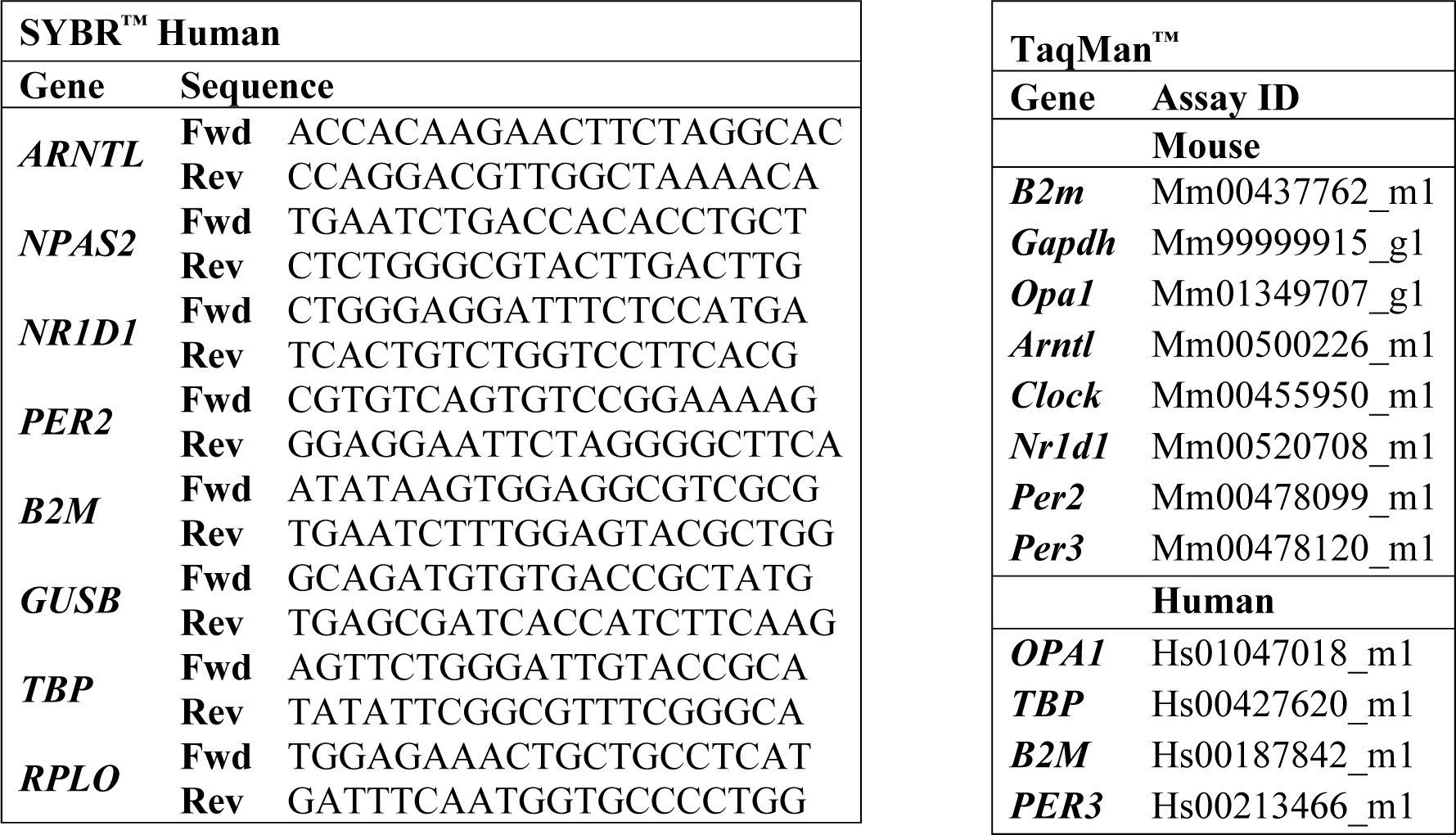
Primer Sequences and Assay IDs used for the Gene Expression Analysis.

#### Immunoblot analysis

Immunoblot analysis was performed as described (Lassiter et al., 2018) with antibodies against OPA1 (#80471) from Cell Signaling Technology, Total OXPHOS (#110411) from Abcam (UK) and β-actin (#A5441) from Sigma-Aldrich. Protein abundance was normalized to β-actin.

### Live-cell microscopy and determination of mitochondrial ROS

Primary human skeletal muscle cells from men with NGT were used for the determination of mitochondrial ROS by live-cell microscopy. Primary skeletal muscle cells were grown and differentiated as described above. MitoSOX (50 µg, Thermo Fisher: M36008) was dissolved in 50 µl pluoronic F127 (Invitrogen: P6866) and 50 µl DMSO to make the MitoSOX stock solution. MitoSOX stock solution (5 µl) was added to the media and incubated for 15 min at 37°C and 5% CO2. Thereafter, media from the cell dish were removed, the dish was mounted on the confocal microscope (Zeiss LSM710) and perfused with Tyrode solution (in mM): 121 NaCl, 5 KCl, 1.8 CaCl2, 0.4 NaH2PO4, 0.5 MgCl2, 24 NaHCO3, 0.1 EDTA, and 5.5 glucose. Cells were then imaged with 2 mins interval for 20 min and thereafter switched to Tyrode solution supplemented with FCCP (2 µM) for 20 min. All images were analyzed with Image J.

---

**Supplementary Figure 1.**
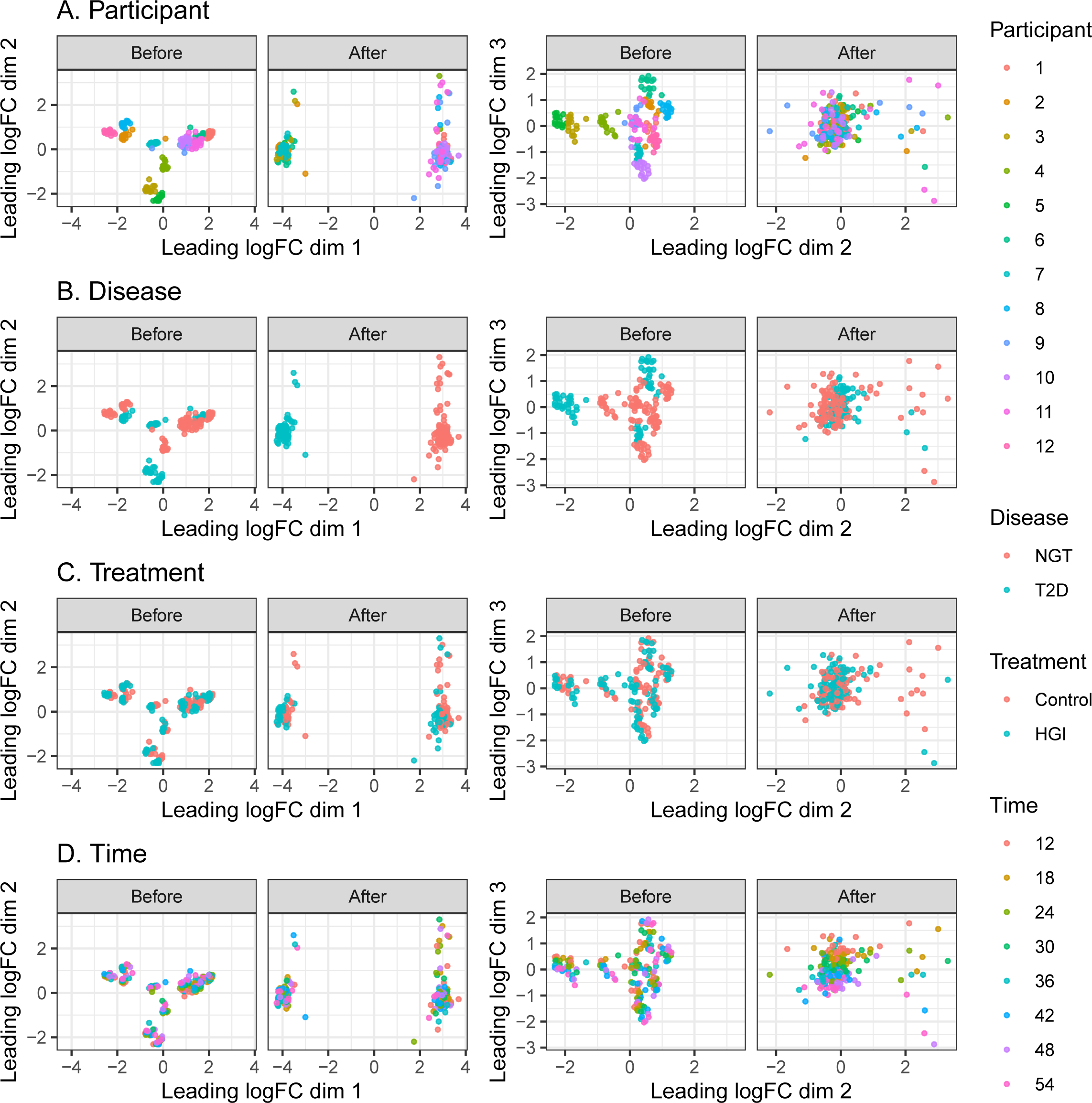
Multi-dimensional scaling (MDS) plots showing dimensions 1-2 (left panels) and 2-3 (right panels) before and after patient batch effect removal. Each experimental condition is presented in A) Patient, B) Disease, C) Treatment and D) Time. Dimension 1 separates the disease group while dimension 3 separates time of sampling after removing the patient batch effect. Time is ZT hours.

**Supplemental Figure S2.**
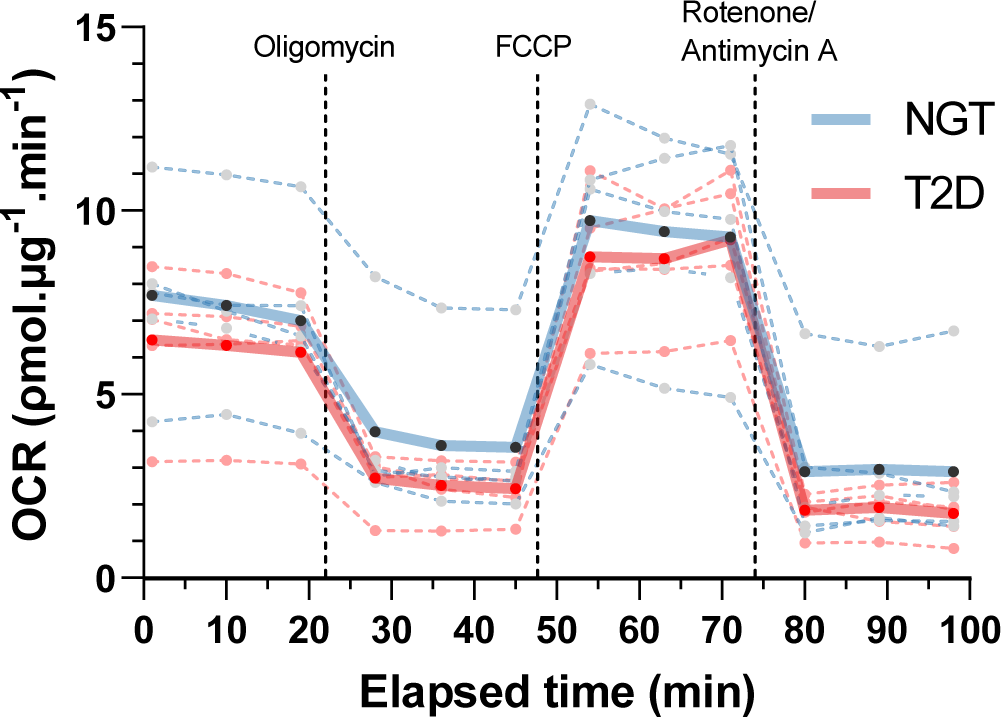
Mitochondrial respiratory function (Mito Stress test, Seahorse XF analyzer) at ZT24 in synchronized myotubes from donors with normal glucose tolerance (NGT; n=5) or type 2 diabetes (T2D; n=5). 2-way repeated measures ANOVA of time-course. Solid lines are overall mean values, while dashed lines are means of individual donors.

**Supplementary Figure 3.**
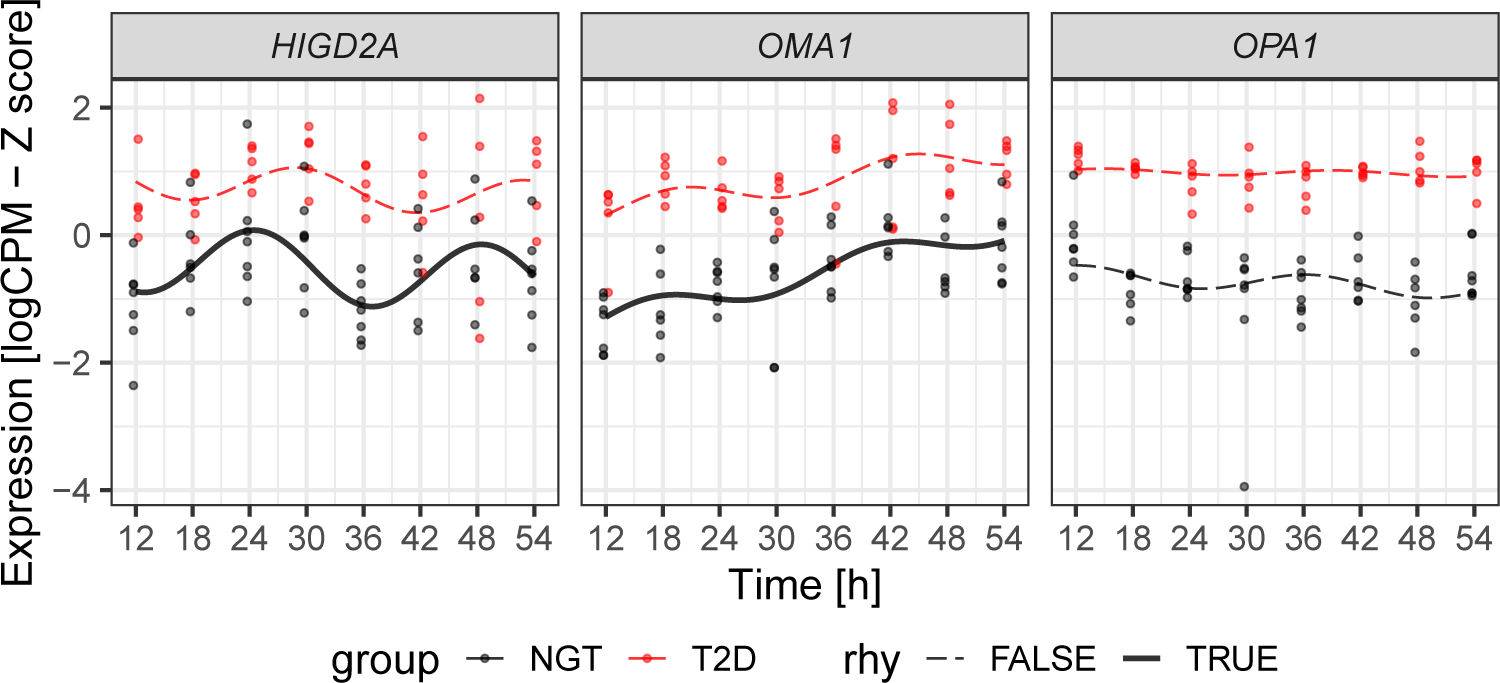
mRNA expression data from RNA sequencing of myotube cultures from donors with NGT or T2D (n=7, and 5, respectively) of *HIGD2A*, *OMA1*, and *OPA1*. Lines show the harmonic regression fits and solid line indicates circadian (adj.*p*<0.1) genes while dashed lines indicate non-circadian genes (RAIN analysis). Time points are hours post-synchronization. Red=T2D, Black=NGT.

Supplemental Table 1.

Rhythmicity analysis results. Each group A) NGT (Control), B) T2D (Control), C) NGT (high glucose and insulin condition) and D) T2D (high glucose and insulin condition) was analyzed with RAIN using longitudinal method. Columns: Group, analysis group; ENSEMBL ID, gene ID from ENSEMBL database; Gene, gene name/symbol; phase and peaks.shape, phase & peak shape estimates from RAIN; period, period of rhythmicity (24 hours); pVal, p-value; pVal.adj, p-values adjusted for multiple comparison by Benjamini-Hochberg method; peak, peak time (hours); trough, trough time (hours); amplitude and phase.estimate; amplitude and phase (peak time) calculated with mean centered data using harmonic regression; description, information about gene; chromosome-start-end-strand, genomic location of the gene.

Supplemental Table 2.

Differential rhythmicity analysis results. T2D was compared to NGT in A) Control and B) high concentration of glucose and insulin conditions. The analysis was performed using DODR algorithm. Columns: Contrast, comparisons made; ENSEMBL ID, gene ID from ENSEMBL database; Gene, gene name/symbol; robustDODR_pVal/ pVal.adj, raw/adjusted p-values from robustDODR method; robustHarmScaleTest_pVal/ pVal.adj, raw/adjusted p-values from robustHarmScaleTest method; meta_pVal/ pVal.adj, combined (meta) raw/adjusted p-values from robust (robustDODR and robustHarmScaleTest); log2FC(amplitude), log2-fold-change of the relative amplitude (T2D vs high concentration of glucose and insulin); description, information about gene; chromosome-start-end-strand, genomic location of the gene

Supplemental Table 3.

Amplitude and phase estimates were calculated using harmonic regression. Relative amplitudes and phases (peak times) of A) NGT (Control), B) T2D (Control), C) NGT (High glucose and insulin) and D) T2D (high glucose and insulin) groups. Log2-ratio (T2D vs. NGT) of relative amplitudes and phase differences are presented for E) Control and F) high glucose and insulin treatment group. The analysis was performed using HarmonicRegression R package. Columns: Group, disease-treatment group; ENSEMBL ID, gene ID from ENSEMBL database; Gene, gene name/symbol; hr.amplitude/hr.phase/hr.pVal/hr.qVal, relative amplitude/phase (peak time)/p-value/q-value calculated by harmonic regression; log2(Amplitude), log2-fold change of the relative amplitude; delta(phase), phase difference between groups; description, information about gene; chromosome-start-end-strand, genomic location of the gene.

